# Whole-cell multi-target single-molecule super-resolution imaging in 3D with microfluidics and a single-objective tilted light sheet

**DOI:** 10.1101/2023.09.27.559876

**Authors:** Nahima Saliba, Gabriella Gagliano, Anna-Karin Gustavsson

## Abstract

Multi-target single-molecule super-resolution fluorescence microscopy offers a powerful means of understanding the distributions and interplay between multiple subcellular structures at the nanoscale. However, single-molecule super-resolution imaging of whole mammalian cells is often hampered by high fluorescence background and slow acquisition speeds, especially when imaging multiple targets in 3D. In this work, we have mitigated these issues by developing a steerable, dithered, single-objective tilted light sheet for optical sectioning to reduce fluorescence background and a pipeline for 3D nanoprinting microfluidic systems for reflection of the light sheet into the sample. This easily adaptable novel microfluidic fabrication pipeline allows for the incorporation of reflective optics into microfluidic channels without disrupting efficient and automated solution exchange. By combining these innovations with point spread function engineering for nanoscale localization of individual molecules in 3D, deep learning for analysis of overlapping emitters, active 3D stabilization for drift correction and long-term imaging, and Exchange-PAINT for sequential multi-target imaging without chromatic offsets, we demonstrate whole-cell multi-target 3D single-molecule super-resolution imaging with improved precision and imaging speed.

## Introduction

Single-molecule localization microscopy (SMLM) has become an invaluable tool for subcellular investigation at the nanoscale^1–6^. In single-molecule super-resolution (SR) imaging^7^, individual fluorescent molecules are temporally separated and localized, thereby surpassing the resolution limit set by the diffraction of light. Exchange-PAINT^8^, i.e. sequential DNA points accumulation for imaging in nanoscale topography (DNA-PAINT), is a localization-based SR approach that involves labeling multiple targets of interest with antibodies conjugated with oligonucleotide strands, called docking strands, and imaging the targets sequentially by introducing the complementary oligonucleotide-dye conjugated sequences, called imager strands. As multiple structures can be labeled using different oligonucleotide pairs and imaged sequentially using the same fluorophore, offsets caused by chromatic aberrations typical in multi-color approaches^9^ can be avoided. One challenge with Exchange-PAINT is that it requires perfusion of the various imager strand solutions and buffers without disturbing the image acquisition process. Another challenge with this method is the increased fluorescence background from freely diffusing imager strands, which increases the achievable single-molecule localization precision^10^. The requirement of low imager strand concentrations for reduced fluorescence background leads to long acquisition times in order to acquire sufficient localization density to achieve good resolution, especially when imaging multiple targets. Recently developed self-quenching fluorogenic DNA probes^11^ reduce background fluorescence in DNA-PAINT imaging, but suffer from an increased localization precision compared to conventional DNA-PAINT probes due to their short binding times which restrict photon collection.

Here, we present soTILT3D, a platform where we developed a steerable, dithered, single-objective tilted light sheet (LS) and a nanoprinted microfluidic chip and combined them with point spread function (PSF) engineering^7^, deep learning^12^, and active stabilization drift correction^13^ to achieve fast, accurate, and precise 3D single-molecule multi-target SR whole-cell imaging using Exchange-PAINT. The single-objective tilted LS and the microfluidic chip fabrication scheme provide the background reduction and solution exchange necessary to perform high-density, 3D, Exchange-PAINT imaging in thick mammalian cells.

LS illumination^2,7,14–16^, where the sample is illuminated with a thin sheet of light at the image plane, has emerged as a simple and effective method of reducing fluorescence background, phototoxicity, and photobleaching of fluorophores. LS illumination offers distinct benefits over other illumination modalities such as total internal reflection fluorescence (TIRF) or highly inclined and laminated optical sheet (HILO)^17^ and variable-angle epifluorescence microscopy (VAEM)^18^ illumination for imaging throughout thick samples due to its ability to illuminate with a thin LS beam and optically section the sample well above the coverslip. Several LS approaches involving the use of separate objectives for illumination and detection have been designed for single-molecule imaging in cells. However, they suffer from optical complexity, low numerical aperture (NA) objectives, steric hindrance, the inability to optically section adherent cells, incompatibility with microfluidic systems, or drift between the objectives^14,19–30^. Previous single-objective LS designs have been demonstrated to reduce the complexity and limitations of multi-objective systems, but they have been limited by beam thickness^17^, the need for beam scanning^31,32^, and limited effective NA in the detection path and the requirement for post-processing due to an illumination beam that is not aligned to the detection axis^33–37^. Our approach based on a fully steerable single-objective tilted LS together with engineered PSFs solves all issues above and enables the use of a single high-NA objective lens for LS focusing and high photon collection efficiency without steric hindrance or relative drift, optical sectioning of entire adherent cells cultured on regular coverslips, and easy combination with microfluidic chips while avoiding the LS aberrations that occur when imaging through microfluidic chip walls^25,33,38–41^. When imaging above the coverslip, in contrast to other single-objective LS methods^33–37^, soTILT3D enables 2D imaging parallel to the image plane and enables 3D imaging down to the coverslip without the need to rotate the image plane. By using LS illumination for Exchange-PAINT imaging, freely diffusing imager strands outside of the focal plane are not excited, resulting in reduced fluorescence background and improved single-molecule localization precision. Tilting the LS enables cellular imaging all the way down to the coverslip without special sample mounts^15^. In addition, our single-objective LS is dithered to achieve homogeneous illumination and reduce shadowing artifacts^19^.

In this work, we develop an easily customizable microfluidic device that incorporates reflective optics into a perfusable microchannel. Previously developed single-objective LS systems have utilized various reflection mechanisms such as a micromirror^42^, an AFM cantilever mirror^43^, pyramidal microcavities^44^, a glass prism^45^, and a microfluidic chamber with metalized side walls^46^ to redirect the LS into the sample. However, these systems use imaging chambers that do not offer design flexibility^21,42–44,46,47^, active solution exchange^21,42–44,47^, or compatibility with epi-illumination and transmission microscopy^42,46^. Our approach solves these issues with a customizable 3D nanoprinted metalized microfluidic insert in a polydimethylsiloxane (PDMS) channel^48–53^ for a user-specified LS tilt angle and chip dimensions and a transparent top for flexible combination with other imaging modalities. Furthermore, the biocompatibility of all materials used makes it an ideal system for studying biological systems with precise control of the extracellular environment. Beyond design flexibility, our microfluidic channel, through its active perfusion compatibility, eliminates the need for manual exchange of solutions which potentially offsets the field of view being imaged and induces drift, reduces thermal damage to the sample by removing heat via constant flow, and increases the number and quality of localizations for DNA-PAINT applications^45^.

To extend SR imaging to 3D, soTILT3D employs PSF engineering, where the shape of the PSF is modulated to encode axial information about the molecule of interest over a several microns axial range in a single slice. PSF engineering is relatively simple to implement, requiring minimal additions to the emission path^7^. Several different PSFs have been designed for this purpose^54–60^. Here, we use the double-helix PSF^15,61,62^ (DH-PSF), which is robust to aberrations, offers good 3D single-molecule localization precision, and requires relatively low computational costs. In contrast to other 3D single-molecule methods, such as multifocus microscopy, PSF engineering bypasses the need for beam splitters and consequent split of photons in the emission pathway while still offering a relatively thick axial range of several microns^47,63^.

SoTILT3D combines PSF engineering with the deep learning based single-molecule localization software DECODE^12^ to enable localization of overlapping emitters in 3D. Due to the enlarged footprint of engineered PSFs, conventional 3D imaging with engineered PSFs requires low emitter density to ensure that no overlap occurs between PSFs. Combined with the low imager strand concentrations already required to reduce fluorescence background during Exchange-PAINT imaging, whole-cell 3D single-molecule SR imaging with engineered PSFs can be a time-consuming process, often taking multiple days to complete. We drastically improve the speed of sequential imaging with Exchange-PAINT by using high concentrations of imager probes. We are able to successfully do this through the reduction of out-of-focus background fluorescence with LS illumination and the use of a deep learning-based localization analysis to localize overlapping emitters^64–67^. While previous work has used PAINT approaches and deep learning to localize 2D data^67^, our approach incorporates LS illumination for optimized signal-to-background ratio (SBR) and adapts the system for 3D analysis of DH-PSF data.

Furthermore, soTILT3D utilizes an active stabilization feedback loop^13^ to correct for drift during image acquisition and enable long-term imaging.

Taken together, soTILT3D combines a steerable, dithered, single-objective tilted LS with microfluidics, PSF engineering, deep learning, and active stabilization drift correction to create a flexible imaging platform for whole-cell multi-target 3D single-molecule SR imaging through Exchange-PAINT with improved localization precision and imaging speeds. We quantify the background reduction and 3D localization precision improvement by performing 3D single-molecule SR imaging of the nuclear lamina protein lamin B1 with soTILT3D compared to conventional epi-illumination. We quantify the speed improvement by imaging α-tubulin and comparing the number of localizations per length of microtubule over time when analyzing with DECODE compared with a non-neural network-based localization software. We then demonstrate the performance of soTILT3D for multi-target 3D single-molecule whole-cell SR imaging of nuclear lamina proteins, mitochondria, and plasma membrane protein ezrin, and quantify the relative distances between nuclear lamina proteins lamin B1, lamin A/C, and lamina associated protein 2 (LAP2). SoTILT3D achieves up to ten-fold faster whole-cell multi-target 3D SR imaging with Fourier ring correlation (FRC) resolution values as low as <30 nm laterally and <40 nm axially, enabling a range of applications for correlating proteins at the nanoscale. We further demonstrate soTILT3D for single-molecule SR imaging of lamin B1 throughout stem cell aggregates using direct stochastic optical reconstruction microscopy (dSTORM)^55,68^, demonstrating applicability and versatility beyond isolated cell samples.

## Results

### SoTILT3D design and performance

SoTILT3D enables 3D single-molecule imaging by reflecting a LS off of a metalized insert with an angled side wall inside of a microfluidic PDMS chip (Fig. 1a-d). The LS is formed using a cylindrical lens, steered with galvanometric mirrors in the x and y directions (Supplementary Figs. 1, 2a,b), and defocused with a tunable lens (Supplementary Figs. 1, 2c,d). An additional galvanometric mirror is used to dither the LS at a half-angle of 20° at a frequency of 100 Hz to reduce striping and shadowing effects (Supplementary Figs. 1, 2e). The resulting LS is 1.1 μm thick (1/e^2^ beam waist radius) (Fig. 1e, Supplementary Fig. 2f), 74.7 μm wide (1/e^2^ diameter) (Supplementary Fig. 2g), and has a confocal parameter of 18.0 μm (1/e^2^) (Supplementary Fig. 2h), and these parameters can easily be tuned by the choice of lenses (see Methods). The Strehl ratio of our system at 5 µm from the coverslip was determined to be 0.85 (Supplementary Fig. 3). The angle and dimensions of the nanoprinted insert as well as the dimensions of the PDMS chip are easily tunable (Fig. 1a-d). Here, a LS beam angle of 12° was used for whole-cell imaging to enable sectioning of adherent cells all the way down to the coverslip. The tilt angle of 12° was chosen to be slightly larger than the ∼10° LS convergence angle, allowing for imaging all the way down to the coverslip without inducing scattering, aberrations, and reflections that would degrade the optical sectioning performance of the LS. The transparent top of the microfluidic chip allows for easy combination with other epi– and transmission illumination modalities. A 4f-system was implemented in the emission path to access the Fourier plane for PSF engineering with a DH phase mask (Fig. 1f, Supplementary Fig. 4) to enable 3D localization of single molecules.

**Fig 1.**
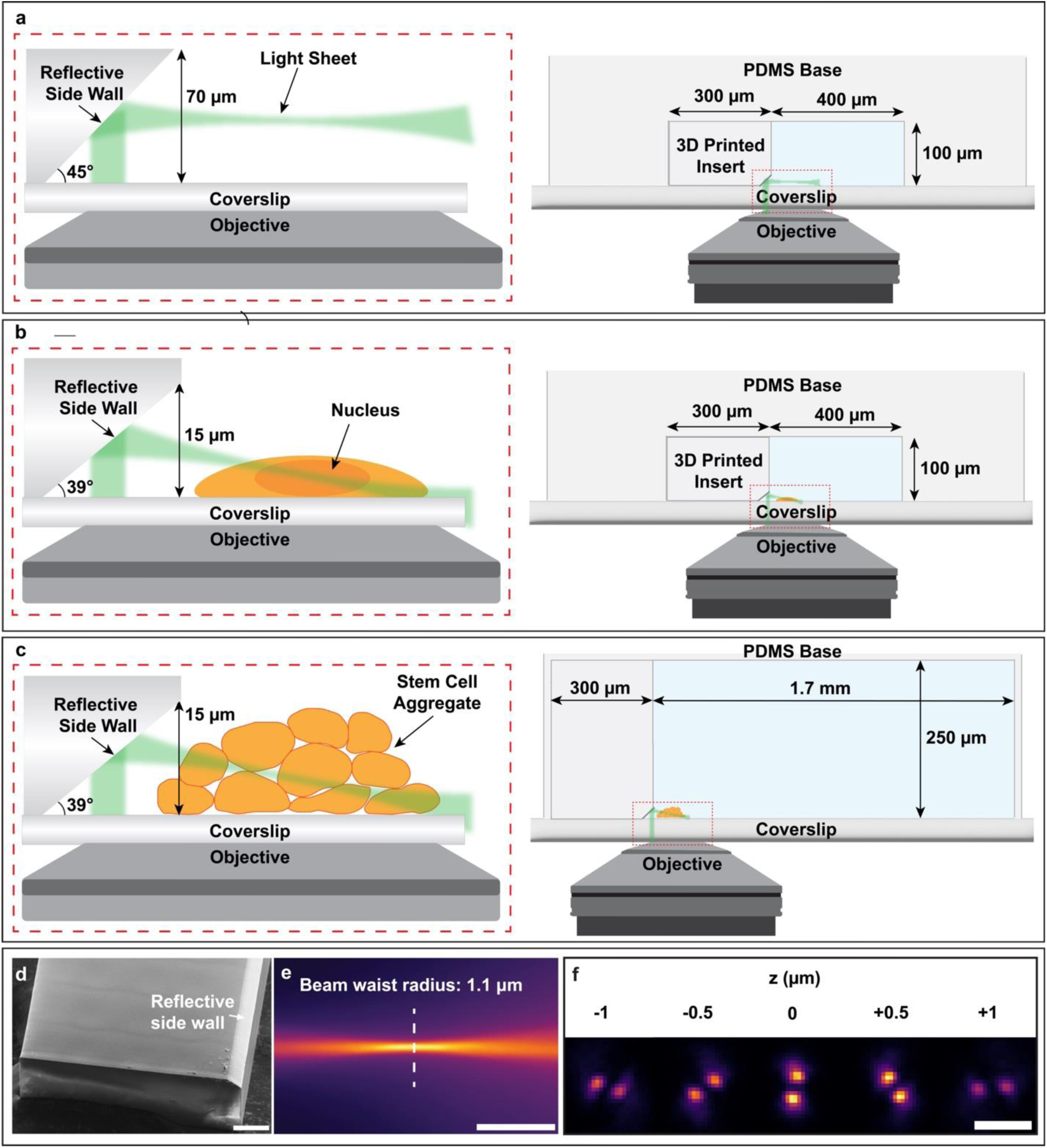
Illustration and characterization of the soTILT3D approach. **a-c** SoTILT3D combines a microfluidic chip that has a nanoprinted insert with metalized side wall with a single-objective tilted light sheet (LS) and is compatible with epi– and transmission illumination. The geometry and size of the insert and the microfluidic chip can be flexibly adjusted for different targets. Schematics not to scale. **d** Scanning electron micrograph of the microfluidic chip insert design shown in b. Scale bar 50 μm. **e** The thin end of the LS imaged in a fluorescent solution. Scale bar 10 μm. **f** Fluorescent beads imaged with the 2 μm axial range double helix point spread function (DH-PSF) at the indicated positions. Scale bar 2 μm.

### SoTILT3D improves the localization precision of 3D single-molecule imaging

Using the LS improves the SBR over 4x for diffraction-limited imaging (Fig. 2a, Supplementary Fig. 5) and over 6x for single-molecule imaging (Fig. 2b), as demonstrated for imaging of lamin B1 in U2OS cells. Single-molecule DNA-PAINT data of lamin B1 in U2OS cells was acquired in 3D using the DH-PSF with epi-or LS illumination to assess the performance of the soTILT3D platform in terms of localization precision (Fig. 2c, Supplementary Fig. 6). In a representative example (Fig. 2c), while roughly matching the signal intensities from the individual molecules with median values of 8,400 photons/localization for epi– and 11,400 photons/localization for LS illumination, the fluorescence background was drastically reduced when using LS illumination and resulted in median background values of 125 photons/pixel for epi-compared to 64 photons/pixel for LS illumination (Fig. 2c). This resulted in improvements in lateral and axial localization precisions, with median values for epi-illumination of 11.2 nm in xy and 16.9 in z and median values for LS illumination of 7.8 nm in xy and 11.8 nm in z (Fig. 2c). The median lateral and axial localization precisions across three technical replicates were found to be 17.9 nm, 14.7 nm, and 10.7 nm in xy and 26.8 nm, 22.1 nm, and 16.1 in z for epi-illumination, while the median values for LS illumination were 9.8 nm, 7.8 nm, and 7.4 nm in xy and 11.9 nm, 11.9 nm, and 11.3 nm in z (Supplementary Fig. 6).

**Fig 2.**
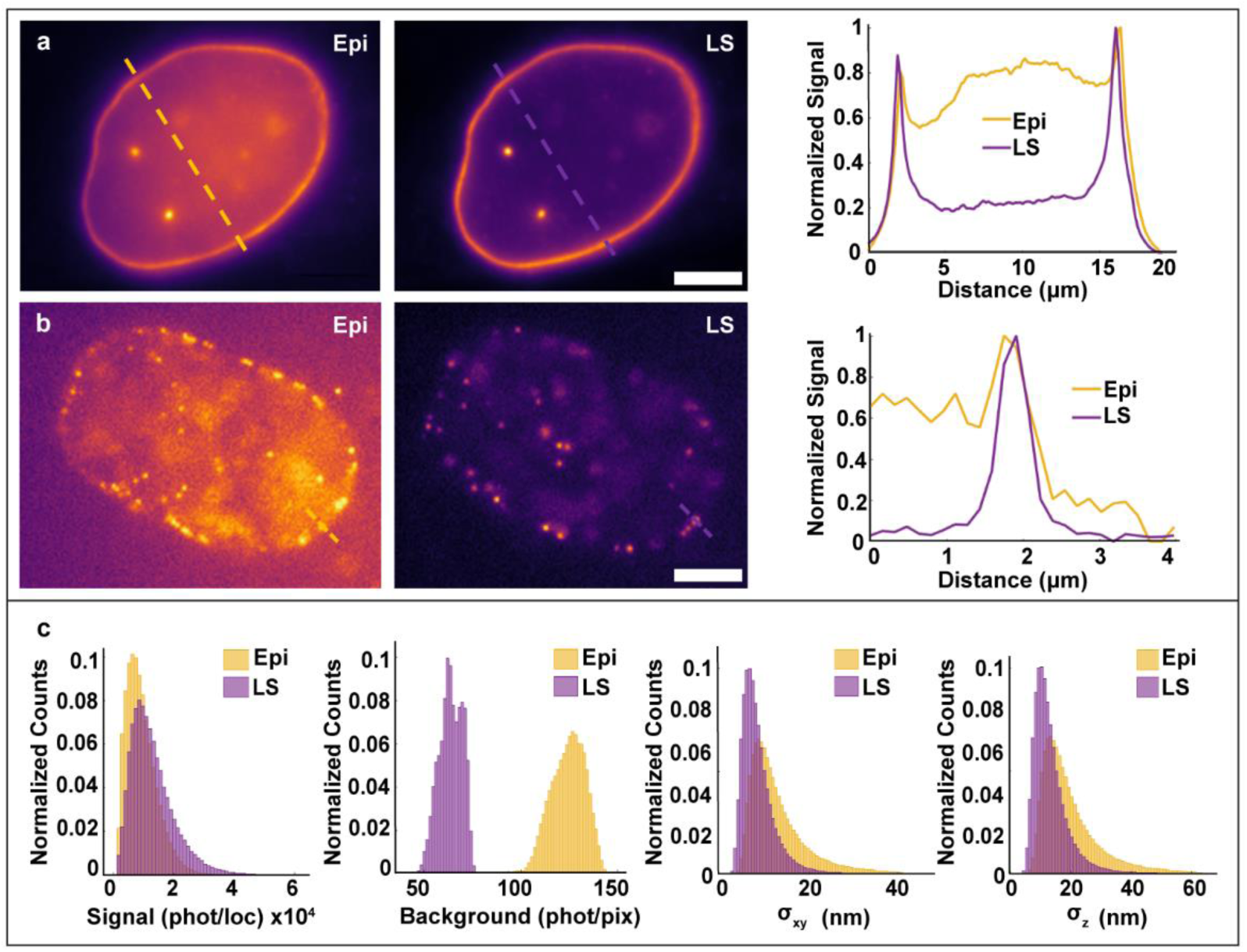
SoTILT3D offers improved signal-to-background ratio and 3D localization precision compared with conventional widefield epi-illumination. **a** Diffraction-limited and **b** single-molecule images of lamin B1 in U2OS cells excited with epi-or light sheet (LS) illumination. Graphs show line scans demonstrating the contrast improvement when using LS compared to epi-illumination. Scale bars 5 μm. **c** Histograms demonstrating comparable signal photon levels but decreased fluorescence background levels leading to improved localization precisions in xy and z for LS compared to epi-illumination for 3D single-molecule super-resolution imaging of lamin B1.

### SoTILT3D improves the speed of 3D single-molecule imaging

The speed improvement of soTILT3D together with deep learning analysis was quantified by 3D single-molecule SR imaging of microtubules in U2OS cells.

First, the approach was benchmarked by analysis of the same fluorescent bead data and single-molecule data with Easy-DHPSF^69,70^, an open-source least-squares fitting-based analysis software, and DECODE, which resulted in comparable localization precisions (Supplementary Fig. 7).

Then, the density breaking point at which 3D single-molecule data could no longer be localized in Easy-DHPSF due to overlapping emitters and at which analysis with DECODE becomes necessary was quantified (Supplementary Fig. 8). The number of detected localizations with Easy-DHPSF was found to decrease after 0.025 nM as the emitter concentration increased (Supplementary Fig. 8a,b). The high-density data was localized with a DECODE model which was trained on the 0.025 nM microtubule data set and converged to a Jaccard Index of 0.69 after 1000 epochs. After 25,000 frames at an imager strand concentration of 0.2 nM, the high-density regime, only 1,222 emitters were localized with Easy-DHPSF compared to 108,254 localized with DECODE (Supplementary Fig. 8c,d). The resulting Easy-DHPSF resolution values were also found to be degraded in comparison to those in the DECODE case (Supplementary Fig. 8c,d).

Next, high density data was acquired with a 0.2 nM concentration of imager strands while our standard concentration data for non-overlapping emitters was acquired at 0.025 nM (Fig. 3a,b). The non-overlapping single-molecule data was localized with Easy-DHPSF and the high-density data was localized with DECODE. This resulted in a more than ten-fold increase in possible imaging speed when using DECODE in conjunction with LS illumination (Fig. 3a,b). The achieved resolutions after 50,000 frames were estimated using FRC calculations in the xy, yz, and xz planes, resulting in resolutions of 35.4/38.6/36.9 nm and 28.4/32.1/33.0 nm for the Easy-DHPSF data set and the DECODE data set, respectively, in the representative example shown in Fig. 3c,d, demonstrating overall improved resolution when using DECODE for the same number of frames. The mean resolution across three technical replicates (Supplementary Fig. 9) was 38.9 ± 2.1 nm, 35.5 ± 2.0 nm, and 32.5 ± 1.2 nm for the Easy-DHPSF data sets and 25.5 ± 0.7 nm, 29.1 ± 0.5 nm, and 27.6 ± 0.2 nm for the DECODE data sets (reported as mean ± standard error of the mean) in the xy, xz, and yz planes, respectively.

**Fig 3.**
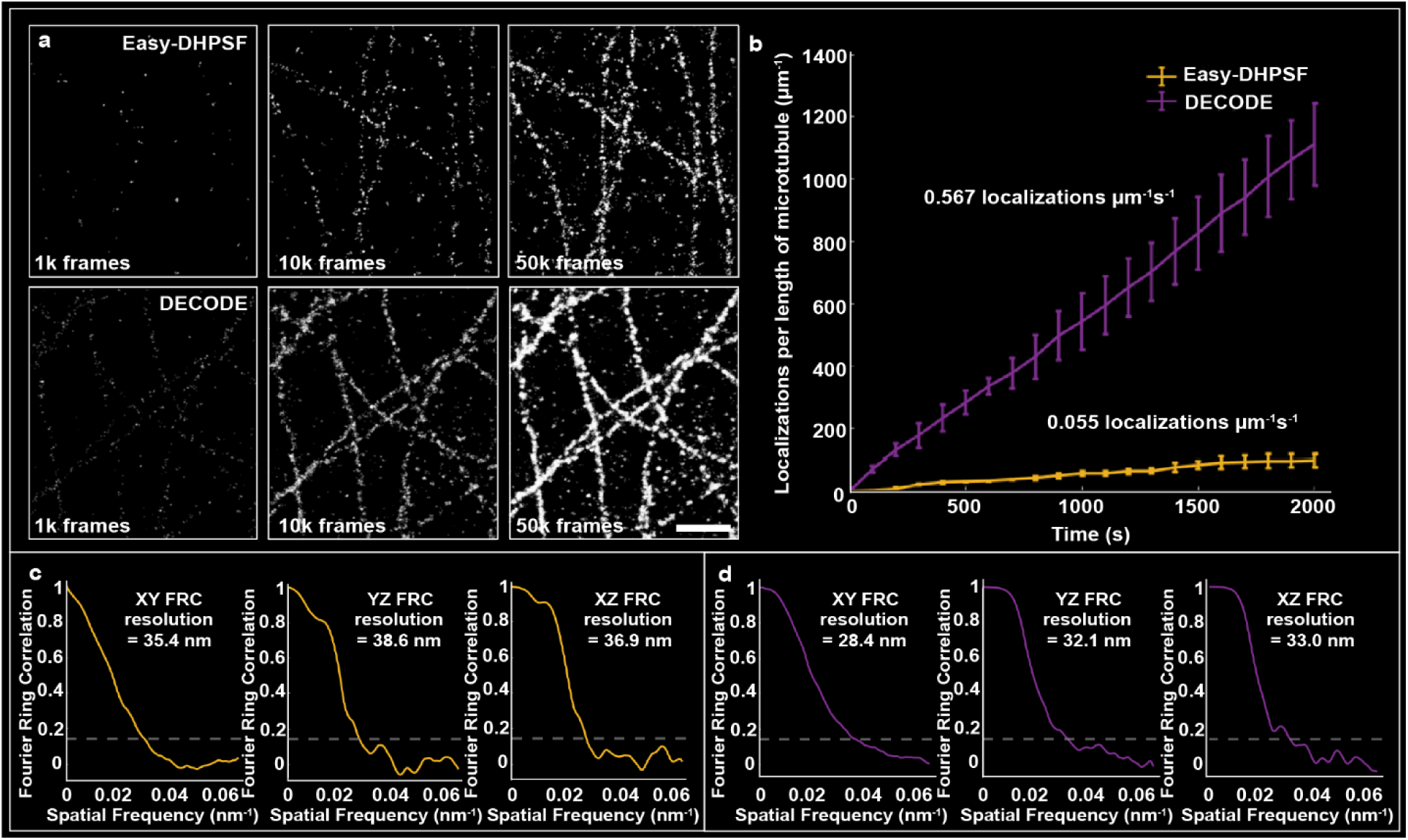
SoTILT3D demonstrates increased 3D acquisition speeds compared with conventional analysis methods. **a** Projections of 3D super-resolution reconstructions of microtubules after acquiring 1,000, 10,000, and 50,000 image frames with 0.025 nM imager strands and analyzed with least-squares fitting-based analysis software (Easy-DHPSF, top row) or with 0.2 nM imager strands and analyzed using a deep learning based approach (DECODE, bottom row). Scale bar 1 μm. **b** Quantitative comparison of the number of localizations per length of microtubule over time achieved with Easy-DHPSF (yellow) and DECODE (purple). The average localizations per µm per second are shown in text above each line, demonstrating a ten-fold increase in speed when using DECODE. Each data point on the graph represents the average localizations per length of microtubule for three different microtubule sections and error bars are ± standard deviations of these data sets. **c** Fourier ring correlation (FRC) curves in the xy, yz, and xz planes after 50,000 frames for the Easy-DHPSF-analyzed data and **d** the DECODE-analyzed data for the representative example shown in a, demonstrating improved resolution when using DECODE.

Finally, 3D high-density single-molecule super-resolution imaging of three separate cells labeled for lamin A/C revealed an increase in the number of localizations detected by DECODE and thus in the acquisition speed, as well as an improvement in the resolution when using LS as the illumination modality compared to epi-illumination (Supplementary Fig. 10). The mean resolution for three technical replicates in the xy, xz, and yz planes when imaged with epi-illumination was 41.9 ± 0.4 nm, 55.8 ± 3.6 nm, and 50.3 ± 1.8 nm, while the mean resolution when imaged with LS-illumination was 39.9 ± 1.9 nm, 48.9 ± 3.5 nm, and 47.1 ± 1.5 nm (reported as mean ± standard error of the mean, Supplementary Fig. 10a,b). The number of localizations detected for the three technical replicates when imaged with epi-illumination was 233,716, 341,101, and 604,412 for cell 1, cell 2, and cell 3 respectively, while the number of localizations when imaged with LS-illumination was 1,215,297, 660,758, and 637,511 for cell 1, cell 2, and cell 3 respectively (Supplementary Fig. 10a,b). Next, the LS-illuminated data sets were filtered for localization precision to match the number of localizations acquired in the epi-illuminated data sets (Supplementary Fig. 10c). The resulting resolution values in the xy, xz, and yz planes were 35.1 ± 3.0 nm, 42.5 ± 4.0 nm, 40.7 ± 4.2 nm (reported as mean ± standard error of the mean) for cell 1, cell 2, and cell 3 respectively. This demonstrates the necessity of the soTILT3D approach to enable imaging in the high-density regime at increased speeds and with improved resolution.

### Whole-cell multi-target 3D single-molecule super-resolution imaging with soTILT3D

Whole-cell single-molecule data was acquired using soTILT3D and DECODE for analysis to demonstrate the performance of soTILT3D for accurate and precise multi-target 3D SR imaging (Figs. 4, 5). The focal plane and LS were moved in 1 µm steps to image each target in overlapping slices while a pressure-driven pump was used to introduce imager strand sequences sequentially.

**Fig. 4.**
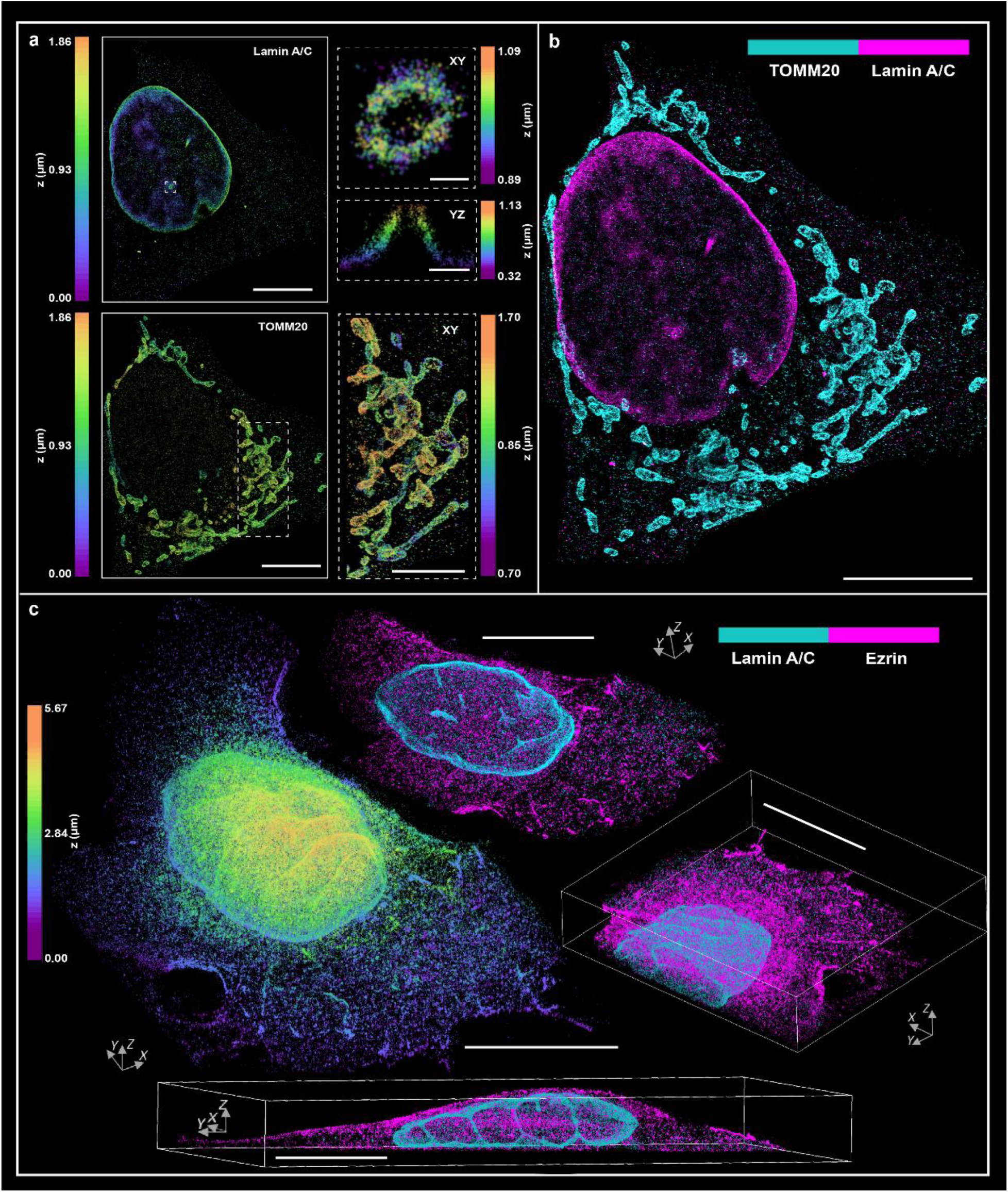
Two-target 3D high-density whole-cell single-molecule super-resolution (SR) imaging with soTILT3D. **a** 3D SR reconstruction of separately rendered lamin A/C and mitochondria (TOMM20) in the same U2OS cell, revealing no notable crosstalk between the two targets. Scale bars 10 μm. Inset panels for lamin A/C shown for a 200 nm thick z-slice in the xy plane and for a 150 nm thick x-slice in the yz plane resolved the nanoscale architecture of a nuclear lamina channel. Xy inset scale bar 200 nm and yz inset scale bar 500 nm. Inset panel for mitochondria shows details of the mitochondrial network. Mitochondrial inset scale bar 5 μm. **b** A merge of the 3D SR reconstructions of lamin A/C and mitochondria shown in a. Scale bar 10 μm. **c** 3D whole-cell SR reconstruction of lamin A/C and ezrin. The left panel shows a merged reconstruction of both targets, colored by depth. The top panel shows a 1.2 μm thick z-slice colored by probe. The right panel shows the bottom right half of the reconstruction in the left panel. The bottom panel shows a 2.5 μm thick slice through the center of the cell. Scale bars 10 μm.

First, 3D two-target imaging was demonstrated by imaging of lamin A/C and mitochondria in U2OS cells, clearly resolving nuclear lamina channels and the intricate mitochondrial network (Fig. 4a,b). The resulting FRC resolutions in the xy/yz/xz planes were found to be 35.0/45.7/51.5 nm for lamin A/C and 32.7/40.2/44.9 nm for mitochondria (Supplementary Fig. 11).

Next, 3D two-target imaging of whole cells was demonstrated by imaging of membrane-bound protein ezrin and lamin A/C in U2OS cells, where the entire axial profile of the cell was reconstructed (Fig. 4c). The resulting FRC resolutions in the xy/yz/xz planes were found to be 31.0/39.2/38.2 nm for ezrin and 38.0/46.3/47.2 nm for lamin A/C (Supplementary Fig. 11).

Next, whole-cell multi-target imaging was demonstrated on nuclear proteins lamin B1, LAP2, and lamin A/C in U2OS cells (Fig. 5). The nuclear protein distributions were resolved with only 10,000 frames per slice (Fig. 5a), as high-density data analyzed with DECODE allows for a greater number of localizations in a shorter time. The nanoscale separation between these nuclear targets were then quantified from these 3D SR reconstructions using Gaussian fits of line scans across the nuclear rim and found to be 41 ± 8 nm for lamin B1 and lamin A/C, 29 ± 9 nm for lamin B1 and LAP2, and 12 ± 10 nm for LAP2 and lamin A/C (reported as mean ± standard error of the mean for 25 line scans) (Fig. 5b-d and Supplementary Fig. 12). Representative fits to a line scan can be seen in Fig. 5c, where nuclear protein distribution distances were found to be 41 nm between lamin B1 and lamin A/C, 27 nm between lamin B1 and LAP2, and 14 nm between LAP2 and lamin A/C. Average nuclear target distributions across all line scans are shown pairwise with Gaussian fits to the average distributions (Fig. 5d). SoTILT3D maintained high resolution throughout the cell during whole-cell, multi-target, 3D SR imaging, as demonstrated by FRC analysis (Fig. 5e, Supplementary Fig. 11). FRC analysis resulted in resolutions in the xy/yz/xz plane of 29.2/36.2/38.6 nm for the whole cell reconstruction and of 27.9/36.1/39.9 nm, 30.2/37.6/38.0 nm, and 31.3/38.9/40.6 nm for lamin B1, LAP2, and lamin A/C, respectively.

**Fig. 5.**
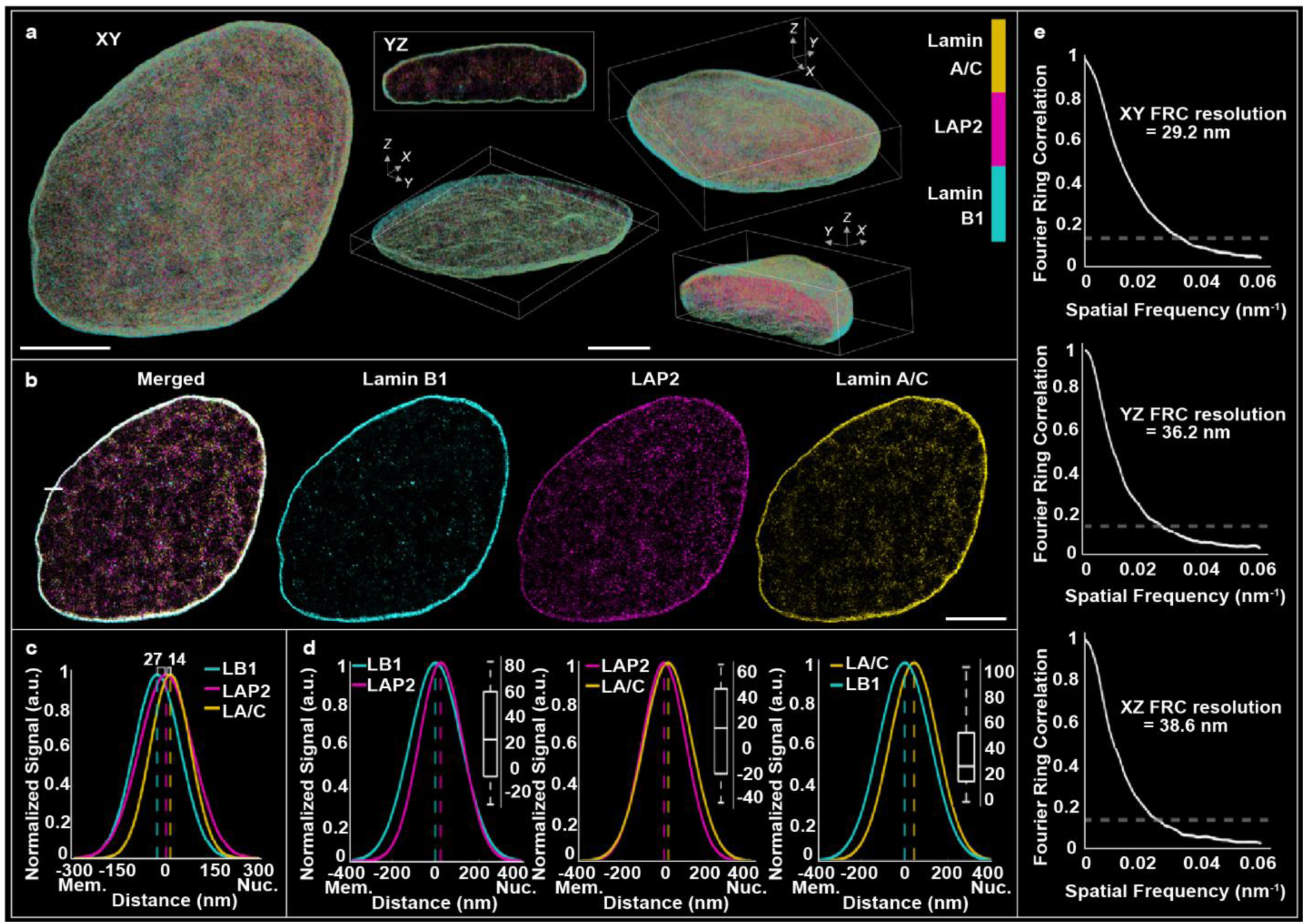
Whole-cell multi-target 3D high-density single-molecule super-resolution (SR) imaging with soTILT3D. **a** 3D multi-target SR reconstruction of an entire U2OS cell nucleus labeled for lamin B1, LAP2, and lamin A/C shown in the xy plane (left), as a 1 μm thick x-slice shown in the yz plane (top, middle), as a 1.5 μm thick z-slice (bottom, middle), at an angle (top, right), and for the cap shown at the bottom left of the left panel (bottom, right). Scale bars 5 μm. **b** 500 nm thick z-slice of the nuclear proteins in a shown in the xy plane merged, and then individually for each target. Scale bar 5 μm. **c** Gaussian fits to an example line scan across the nuclear rim (white line in the merged sample in b), demonstrating the separation distances between lamin B1 (LB1), LAP2, and lamin A/C (LA/C) in nm from the nuclear membrane (Mem.) towards the center of the nucleus (Nuc.). **d** Average intensity profile plots corresponding to line scans across the nuclear rim characterizing distances between lamin B1 and LAP2, LAP2 and lamin A/C, and lamin B1 and lamin A/C. Insert box-and-whisker plots represent the distribution of separations (nm) across all line scans. **e** Fourier ring correlation (FRC) curves in the xy, yz, and xz planes of all three targets in a.

Taken together, the resolution from the FRC analyses were found to be 32.3 ± 1.2 nm, 40.6 ± 1.4 nm, and 42.9 ± 1.8 nm (reported as mean ± standard error of the mean for seven structures, see Supplementary Fig. 11) in the xy, yz, xz plane, respectively, for all targets, demonstrating robust performance across multiple different cellular structures.

Finally, super-resolution imaging of a broader array of biological samples was demonstrated by imaging of lamin B1 in stem cell aggregates using dSTORM (Supplementary Fig. 13). Line scans of diffraction-limited images acquired using epi– and LS illumination revealed up to a 4x improvement in SBR when using LS illumination (Supplementary Fig. 13a-c) and 2D SR images of lamin B1 resolved details of the nuclear architecture throughout the aggregates (Supplementary Fig. 13d-f).

## Discussion

The combination of a steerable, dithered, single-objective tilted LS with microfluidics, PSF engineering, deep learning, active drift stabilization, and Exchange-PAINT resulted in improved localization precision and imaging speeds for whole-cell multi-target 3D single-molecule SR imaging. In contrast to our earlier tilted LS design that utilized two objectives^15^, soTILT3D employs a single, high-NA objective. This configuration enables the formation of a significantly thinner LS, making it well-suited for challenging samples that exhibit higher background fluorescence. The optical system is simple and requires only standard, commercially available optical elements, and can be easily implemented on conventional microscopes.

The developed microfabrication pipeline provides a simple procedure for fabricating microfluidic devices that are compatible with single-objective LS illumination, conventional widefield epi-illumination, and transmission microscopy. The flexibility of the 3D nanoprinted insert geometry along with the wide range of design options for the PDMS channels allow for easy adaptability and implementation for various applications, as demonstrated by imaging of both isolated cells and stem cell aggregates in microchannels of different dimensions. Furthermore, we think our microfluidic fabrication pipeline provides an easy and flexible method for the incorporation of 3D nanoprinted optics in microchannels that paves the way for further studies incorporating microoptics into microfluidic systems.

Transmission-based imaging of polystyrene beads was used for real-time stabilization to keep the sample within the field of view and within the same LS slice during long-term imaging. This approach is based on an open-source plugin for easy implementation^13^. The fluorescent fiducial beads were then used to correct any residual drift in post-processing. An alternative approach is using real-time localization of the fluorescent bead together with real-time feedback to the stage for active stabilization during imaging^15^.

The soTILT3D platform can be easily extended to live-cell imaging and single-particle tracking (SPT). The biocompatibility of the insert and PDMS together with the gas permeability of PDMS makes the microfluidic channels suitable for live-cell studies. In fact, the cells in this study were cultured inside the chip before fixation. Solution perfusion through the channel allows for precise control of the extracellular environment and swift removal of heat or waste products with a constantly replenishing fluid reservoir. The single-objective LS design is also compatible with standard microscope stage top incubators. Furthermore, LS illumination minimizes photodamage to the specimen and the auto-stabilization drift correction scheme allows for easy decoupling of cell movement from sample drift over long acquisition periods. The design flexibility of the soTILT3D setup also allows for easy implementation with other live-cell compatible single-molecule imaging modalities, such as Peptide-PAINT^71,72^ and SR imaging using SiR dyes^73–75^. New fluorophore developments, including self-quenched DNA-PAINT probes^76^, and tunable lenses with stronger defocusing abilities, will further improve the performance of soTILT3D imaging in the future. Additionally, refractive index mismatch-induced aberrations such as spherical aberrations can be reduced in future studies by the use of a silicone oil immersion detection objective for better matching with the refractive index of the sample. Adaptive optics can also be implemented as an addition to the 4f setup to mitigate the effects of setup– and sample-induced aberrations, including spherical aberrations.

Overall, we think that soTILT3D offers a simple and flexible approach for improved 3D SR imaging and SPT that can be adapted and utilized for various single-molecule imaging applications for faster and more efficient and precise nanoscale investigation of cellular structures and molecular dynamics.

## Methods

### Optical platform

The optical platform was built around a conventional inverted microscope (IX83, Olympus) (Supplementary Fig. 1). The excitation pathway includes illumination lasers (560 nm, 1000 mW, MPB Communications; 642 nm, 1000 mW, MPB Communications) which were spectrally filtered (FF01-554/23-25, Semrock; FF01-631/36-25, Semrock) and circularly polarized (560 nm: Z-10-A-.250-B-556 quarter-wave plate, Tower Optical; 642 nm: LPVISC050-MP2 polarizer, Thorlabs, and Z-10-A-.250-B-647 quarter-wave plate, Tower Optical) before being expanded and collimated (LA1951-A, *f* = 25.4 mm, Thorlabs; LA1417-A, *f* = 150 mm, Thorlabs). Switching between lasers was controlled with shutters (VS14S2Z1, Vincent Associates Uniblitz) connected to a shutter control box (VDM-D3, Vincent Associates Uniblitz). The laser pathways were then merged into a single optical path with a dichroic mirror (T5901pxr-UF2, Chroma). The laser light was then either sent to the epi-illumination or single-objective LS illumination pathway, where switching between paths was achieved by manually flipping two flip mirrors. Along the epi-illumination pathway, the beam was further expanded and collimated (AC508-075-A, *f* = 75 mm, Thorlabs; AC508-300-A, *f* = 300 mm, Thorlabs) to achieve a laser spot size that would illuminate the entire field of view (FOV) of ∼50 μm x 50 μm. The beam was reflected off of a flip mirror and directed through a Köhler lens (AC508-300-A, *f* = 300 mm, Thorlabs) that focused the beam at the back aperture of a high-NA oil immersion objective (UPLXAPO100x, 100x, NA 1.45, Olympus) used for both illumination and detection to generate collimated widefield epi-illumination in the sample plane. Along the LS illumination pathway, the beam was reduced and collimated (AC254-15-A-ML, *f* = 150 mm, Thorlabs; AC254-075-A-ML, *f* = 75 mm, Thorlabs) before going through a cylindrical lens (ACY254-150-A, *f* = 150 mm, Thorlabs) which focused the light in only one dimension onto a galvanometric mirror oriented horizontally and conjugated to the back focal plane of the objective lens for beam steering in the sample plane in the y direction (GVS211, Thorlabs; GPS011-US Galvo Power Supply, Thorlabs). The beam was then reduced and collimated once again (AC254-150-A-ML, *f* = 150 mm, Thorlabs; AC254-075-A-ML, *f* = 75 mm, Thorlabs) and focused onto a second galvanometric mirror oriented vertically and conjugated to the back focal plane of the objective lens for beam steering in the sample plane in the x direction (GVS211, Thorlabs; GPS011-US Galvo Power Supply, Thorlabs). The beam was then reduced and collimated once again (AC254-150-A-ML, *f* = 150 mm, Thorlabs; AC254-075-A-ML, *f* = 75 mm, Thorlabs) and focused in one dimension onto a tunable lens (EL-3-10-VIS-26D-FPS, Optotune; EL-E-4 Electrical Lens Driver, Optotune) conjugated to the back focal plane of the objective lens for focal steering of up to 50 μm in the sample plane. The beam was then directed through a lens (AC508-300-A, *f* = 300 mm, Thorlabs) focusing the beam onto a third galvanometric mirror (SP30Y-AG, Thorlabs; GPWR15 Galvo Power Supply, Thorlabs; CBLS3F Galvo System Cable Set, Thorlabs) oriented horizontally and conjugated to the sample plane for rapid dithering of the LS with a function generator (15 MHz DDS Signal Generator/Counter, Koolertron). Finally, the beam was directed through the Köhler lens, a common element between both the epi– and the LS illumination paths. All lenses in the illumination path were chosen to generate a LS with dimensions suitable for mammalian cell imaging. The high-NA objective lens then focused the LS into a microfluidic chip bonded to a coverslip (*vide infra*) and was reflected off of its metalized side wall at a 0° or 12° angle into the sample. For whole-cell SR imaging, mirrors on removable magnetic mounts in the red epi-illumination path were used for simultaneous epi-illumination for fiducial bead-based drift correction (Supplementary Fig. 14). A dichroic mirror (69-216, 600 nm Dichroic Shortpass Filter, Edmund Optics) was placed on a sideways-mounted magnetic mount in place of the flip mirror just before the Köhler lens to allow for simultaneous LS illumination with the 560 nm laser and epi-illumination with the 642 nm laser. This allowed for fiducial bead-based drift correction using 0.20 μm 660/680 fiducial beads (F8807, Invitrogen) in both lateral and axial dimensions while acquiring single-molecule data with the single-objective LS.

Real-time active stabilization was achieved using an infrared (IR) LED (BLS-LCS-0850-03-22, Mightex; BLS-SA02-US LED Control Box, Mightex) mounted in place of the microscope’s white light halogen lamp for IR brightfield illumination. An 800 nm dichroic short-pass mirror (14-015, Edmund) was mounted between the objective lens and the Köhler lens to direct transmitted IR light to a separate path where it was focused using a tube lens (ACT508-200-B, *f* = 200 nm, Thorlabs) onto the sensor of a CMOS camera (CS235MU, Thorlabs) aligned in an optical cage system. Optical cleanup filters (FF01-842/56-25, Semrock) were placed between the LED and sample and in front of the CMOS camera to filter stray IR wavelengths in the transmission path. Furthermore, a short-pass cleanup filter (BSP01-785R-25, Semrock) was placed in the fluorescence detection path to remove residual IR wavelengths from the fluorescence emission light. This allowed for active stabilization of the system using the standard PSF of non-fluorescent 3 μm polystyrene beads (C37484, Invitrogen) in both lateral and axial dimensions using an open-source active stabilization ImageJ/Micromanager Plugin^13^ termed Feedback Focus Lock (Supplementary Fig. 14), which relied on a fine adjustment xyz piezoelectric translation stage (OPH-PINANO-XYZ, Physik Instrumente) to correct for drift while acquiring single-molecule data.

The microfluidic chip was mounted in a stage top incubation chamber (P-736-ZR1S/ZR2S, OkoLab) compatible with an xy translation stage (OPH-XYS-O, Physik Instrumente) and a fine adjustment xyz piezoelectric translation stage (OPH-PINANO-XYZ, Physik Instrumente). Solutions were perfused using a pressure-based flow control pump (LU-FEZ-0345, Fluigent, Inc.) connected to compressed air. Switching between solutions was achieved with an 11-port/10-position bidirectional valve connected to multiple solutions (ESSMSW003, Fluigent, Inc.) and locally controlled using a microfluidic flow controller (ELUSEZ, Fluigent, Inc.). Tubing (FEP 1/16” OD, 1/100” ID, Cole-Parmer) was connected from the pump to the microfluidic chip for solution perfusion.

Emission from fluorophores was collected by the same high-NA objective lens (UPLXAPO100x, 100x, NA 1.45, Olympus) used for epi– and LS illumination, spectrally filtered (ZT405/488/561/640rpcV3 3 mm thick dichroic filter set mounted in Chroma BX3 cube, Chroma; ZET642NF, ZET561NF, both Chroma), and focused by the microscope tube lens to form an image at the intermediate image plane (IIP). The light then continued to the first lens of a 4*f* system (AC508-080-AB, *f* = 80 mm, Thorlabs) positioned one focal length away from the IIP. An iris attached to a lens tube on the emission port of the microscope allowed for control of the emission spot size on the camera sensor. A dichroic mirror (T660lpxr-UF3, Chroma) was used to split the emission into two color channels: a “green path” where wavelengths shorter than 660 nm were reflected into and a “red path” where wavelengths longer than 660 nm were transmitted into. A second 4*f* lens (AC508-080-AB, *f* = 80 mm, Thorlabs) was placed in each path two focal lengths away from the first 4*f* lens. One focal length after the first 4*f* lens, the Fourier plane of the microscope was accessible, and the phase of emitted light was modulated to engineer the PSFs to encode axial positions of emitters in each path. This was achieved using a dielectric DH phase mask (DH-1 phase mask, 590 nm with diameter of 2.367 mm, Double-Helix Optics, LLC) in the green channel for single-molecule data collection over a 2-μm axial range and a DH phase mask (DH-12 phase mask, 670 nm with diameter of 2.484 mm, Double-Helix Optics, LLC) in the red channel for bead localizations and drift correction over a 12-μm axial range. Beyond these ranges, the fluorescence signal of the PSFs becomes dimmer and the rotation rates slow down relative to axial position change. After phase modulation with the phase masks, the emitted light in each color channel was focused by the second 4*f* lens in each channel and directed onto opposite corners of the sensor of an EMCCD camera (iXon Ultra 897, Andor, Oxford Instruments).

For fluorescence imaging, diffraction-limited and single-molecule data was acquired with the Andor Solis EMCCD imaging software. A set EM gain of 200 was used, which corresponded to a calibrated EM gain of 182 for the EMCCD camera. The EMCCD camera conversion gain was found experimentally to be 4.41 photoelectrons per A/D count. This camera was operated at a shift speed of 1.7 μm/s and normal vertical clock voltage amplitude. The read-out rate was set to 10 MHz at 16 bits using a preamplifier gain of 3. The calibrated pixel size of the camera was 159 nm/pixel vertically and 157 nm/pixel horizontally. For transmission imaging, the CMOS camera was set to a gain of 20 photoelectrons per A/D count, and the calibrated pixel size was 42.7 nm/pixel.

### Fabrication process for microfluidic chips with reflective angled side walls

An overview of the fabrication process for microfluidic chips can be found in Supplementary Fig. 15. An overview of the various chip dimensions and modifications used throughout this work can be found in Fig. 1. The microfluidic chip consists of two main parts: a 3D nanoprinted metalized insert and a PDMS channel.

For insert fabrication, inserts of desired dimensions were first designed using AutoCAD (Autodesk) and sliced and hatched for 3D nanoprinting in Describe (Nanoscribe GmbH & Co.KG) with the 10x Silicon Shell recipe using a slicing distance of 0.3 μm and a hatching distance of 0.3 μm with a hatching angle of –6° and a hatching angle offset of 0°. The block size (X: 840 μm, Y: 800 μm, Z: 4000 μm), block offset (X: 400 μm, Y: 425 μm, Z: 0 μm), and block shear angle (0°) were adjusted to avoid any stitching lines throughout the printed structure. For imaging in mammalian cells, inserts were printed to be 105 μm tall, 300 μm wide, and 4 mm long. For imaging throughout stem cells, inserts were printed to be 260 μm tall, 300 μm wide, and 4 mm long. In both cases, the side wall had an angle of 39° and a height of 15 μm to allow for a reflected LS with a 12° tilt. For imaging the LS in fluorescent solution, inserts were printed to be 105 μm tall, 300 μm wide, and 2.5 mm long with an angle of 45° and a height of 70 μm to allow for a reflected LS with no tilt for characterization purposes. A mirror with a height of 70 μm was used to allow for a greater mirror surface area in order to image both the width and thickness of the LS by rotating the cylindrical lens. The design was then printed using two-photon polymerization direct laser writing on a fused silica substrate with the Nanoscribe Photonic Professional GT2 (Nanoscribe GmbH & Co.KG). The structure was printed using the 10x objective with IP-Visio resin (Nanoscribe GmbH & Co.KG), a non-cytotoxic, optically transparent, nonfluorescent methacrylate-based resin, with a laser power set to 100% and a scan speed of 30 mm/s. The structure was then briefly immersed in SU8 developer (mr-Dev 600, Kayaku Advanced Materials, Inc.) where a pipette was used to gently dissolve excess unpolymerized resin. The structure was then briefly immersed in isopropyl alcohol (MPX18304, Fisher Scientific) and dried gently with nitrogen gas. The insert was then treated for 15 minutes with an 18 W UV light source (140010, Vacuum UV-Exposure Box, Gie-Tec GmbH). The insert was then mounted on its side for vapor deposition. This orientation ensured that only the edge of the insert containing the side wall was metalized to reduce unnecessary scattering from reflectivity of other portions of the insert. The insert was then mounted in an e-beam vapor deposition chamber where 200 nm of silica was deposited (EVMSIO21-5D, Kurt J. Lesker), followed by 350 nm of aluminum (EVMAL50EXEB, Kurt J. Lesker), and followed by 5 nm of silica (EVMSIO21-5D, Kurt J. Lesker). The first layer of silica provides an insulating layer to the plastic insert to protect it from the laser intensities used for imaging, the layer of aluminum provides a reflective layer for single-objective LS reflection, and the final layer of silica provides a dielectric coating for the mirror surface. These layers were optimized to provide the best heat protection for the thermoplastic base as well as good reflectivity. Scans profiling the surface of the insert side wall were performed using atomic force microscopy (AFM). A 10 μm x 10 μm region of the side wall was scanned (NCHR-10, NanoWorld; NX20, Park Systems) and analyzed (XEI 4.3.4, Park Systems), revealing a root mean square (RMS) roughness of 25.9 nm for the entire 100 μm^2^ scanned region (Supplementary Fig. 15).

For the PDMS base, PDMS (SYLGARD 184 Silicone Elastomer Kit, Dow Inc.) channels were formed from SU8 molds made on silicon wafers (444, University Wafer, Inc.) with SU8-100 photoresist (Y131273, Kayaku Advanced Materials, Inc.). SU8 molds with a 100 μm height for isolated cell imaging and LS characterization were prepared by spin-coating (WS-650Mz-23NPPB, Laurell Technologies) approximately 2 mL of the photoresist onto a silicon wafer first to 500 revolutions per minute (rpm) at 100 rpm/second acceleration, then held at 500 rpm for 10 seconds, then ramped to 3000 rpm at an acceleration of 300 rpm/second and held at this speed for 30 seconds. The molds were then heated on a hot plate at 65°C for 10 minutes followed by 95°C for 30 minutes. A custom film photomask (Micro Lithography Services Ltd) with channels that were 700 µm wide and 1 cm long designed in AutoCAD (Autodesk) was applied to the photoresist-coated silicon wafers which were then exposed to UV light under vacuum (140010, Vacuum UV-Exposure Box, Gie-Tec GmbH) with an 18 W light source for 27 seconds. The molds were then heated again on a hot plate at 65°C for 1 minute followed by 95°C for 10 minutes. The molds were then placed in a beaker with approximately 20 mL of SU8 developer (mr-Dev 600, Kayaku Advanced Materials, Inc.) and gently swirled for 10 minutes to dissolve residual untreated photoresist. All these steps were performed in accordance with the procedure outlined on the technical data sheet for SU8-100 negative epoxy photoresist (https://kayakuam.com/wp-content/uploads/2020/09/KAM-SU-8-50-100-Datasheet-9.3.20-Final.pdf). SU8 molds with a 250 μm height for stem cell imaging were prepared by spin-coating approximately 2 mL of the photoresist onto a silicon wafer first to 500 rpm at 100 rpm/second acceleration, then held at 500 rpm for 10 seconds, then ramped to 1500 rpm at an acceleration of 300 rpm/second and held at this speed for 30 seconds. The molds were then heated on a hot plate at 65°C for 2 hours followed by 95°C for 4 hours. A photomask with channels that were 2 mm wide and 1 cm long was applied to the photoresist-coated silicon wafers which were then exposed to UV light under vacuum with an 18 W light source for 45 seconds. The molds were then heated again on a hot plate at 65°C for 1 minute followed by 95°C for 30 minutes. The molds were then placed in a beaker with approximately 20 mL of SU8 developer and gently swirled for 20 minutes to dissolve residual untreated photoresist. Once molds were made, approximately 25 mL of PDMS was prepared from a two-part system of a 10:1 ratio of elastomer to curing agent and poured over SU8 molds placed in 60 mm petri dishes (FB0875713A, Fisher Scientific). Pins with cut off tips were placed on both ends of the channel prior to pouring the PDMS and were left in place as the PDMS cured to generate holes for tubing. The PDMS was left to cure at room temperature overnight followed by an additional 2 hours at 65°C in an oven. Once the PDMS was cured, the PDMS slab was peeled off the SU8 mold, and the metalized insert, which can be handled with forceps, was carefully placed into the PDMS channel. Care must be taken to ensure the insert is seated in the correct orientation within the PDMS for LS reflection into the sample. For bonding of the microfluidic chip, coverslips (#1.5, 22 mm x 22 mm, Electron Microscopy Sciences) were first cleaned via sonication (Branson Ultrasonics Cleaning Bath, 15-336-120, Fisher Scientific) in acetone (BP2403-4, Fischer Scientific) for 25 minutes followed by plasma treatment (PDC-32G, Harrick Plasma Inc) with argon gas for 25 minutes. A clean coverslip was then placed with the PDMS into the plasma cleaner (PDC-32G, Harrick Plasma Inc) and treated with air plasma for 30 seconds. Immediately after removal from the plasma cleaner, the PDMS and coverslip were pressed together, allowing covalent bonds to form between the two, and bonding the microfluidic chip to form a sealed channel.

### Human embryonic stem cell line

The human embryonic stem cell line used in this study (ESI-017, courtesy of the Warmflash lab at Rice University) was originally purchased from ESIBIO. Karyotype and genomic integrity checks were done by the manufacturer. Pluripotency (OCT4+, SOX2+, NANOG+) was regularly monitored prior to and during the study. The cell line was tested regularly for mycoplasma and found negative.

All experiments were performed with a human embryonic stem cell line that was previously established and is in the NIH registry. Experiments conform to the ISSCR guidelines found at https://www.isscr.org/policy/guidelines-for-stem-cell-research-and-clinical-translation.

### Cell culture

Before plating cells, the microfluidic channel was cleaned with 70% ethanol (BP82031GAL, Fisher Scientific) followed by rinsing with nanopure water, and coated with a 0.001% solution of fibronectin (F0895, Sigma-Aldrich) in phosphate-buffered saline (PBS) (SH3025601, Fisher Scientific). Human osteosarcoma cells (U-2 OS – HTB-96, ATCC) were then seeded into the microfluidic chips 24 hours before fixation and incubated at 37°C and 5% carbon dioxide (Thermo Scientific Heracell 150i CO_2_ Incubator, 51-032-871, Fisher Scientific) in high-glucose Dulbecco’s modified Eagle’s medium (DMEM, Gibco) with 25 mM HEPES and supplemented with 10% (v/v) fetal bovine serum (FBS, Gibco) and 1 mM sodium pyruvate (Gibco). Human pluripotent stem cells (ESI-017, seeded and fixed in the Warmflash lab at Rice University), were seeded into the microfluidic chips 4 hours before fixation and incubated at 37°C and 5% carbon dioxide (Thermo Scientific Heracell 150i CO2 Incubator, 51-032-871, Fisher Scientific) in mTeSR complete cell culture medium (courtesy of the Warmflash lab at Rice University).

### Sample preparation

After culturing the U2OS cells inside the microfluidic chip for 24 hours, or 4 hours in the case of the stem cells, the cells were fixed and immunolabeled inside of the microfluidic chip. For fixation, cells were first washed three times in PBS (SH3025601, Fisher Scientific), where each wash represents a complete fluid exchange in the microfluidic chip, fixed for 20 min in chilled 4% formaldehyde solution made by dissolving paraformaldehyde (PFA, Electron Microscopy Sciences) in PBS, washed once more in PBS, and incubated with 10 mM ammonium chloride (Sigma-Aldrich) in PBS for 10 minutes. The cells were then permeabilized with three washing steps with 0.2% (v/v) Triton X-100 (Sigma-Aldrich) in PBS with a five-minute incubation period between each wash, and blocked with 3% (w/v) bovine serum albumin (BSA, Sigma-Aldrich) in PBS for 1 hour in the case of the U2OS cells, and in 10% (v/v) donkey serum (ab7475, Abcam) in PBS for 1 hour in the case of the stem cells.

For diffraction-limited imaging of U2OS cells, cells were labeled with rabbit anti-lamin B1 (ab16048, Abcam) primary antibodies using a 1:1,000 dilution in 1% (w/v) BSA in PBS for two hours. Cells were then washed three times with 0.1% (v/v) Triton X-100 in PBS with a three-minute incubation period during each wash before being labeled with donkey anti-rabbit secondary antibodies conjugated with dye CF568 (20098-1, Biotium) at a 1:100 dilution in 1% (w/v) BSA in PBS for one hour. Cells were then washed five times in 0.1% (v/v) Triton X-100 in PBS. An oxygen scavenging buffer^77^ containing 100 mM Tris-HCl (J22638-K2, Thermo Scientific), 10% (w/v) glucose (215530, BD Difco), 2 µl/ml catalase (C100, Sigma-Aldrich), and 560 µg/ml glucose oxidase (G2133, Sigma-Aldrich) was added during imaging to reduce photobleaching.

For single-molecule imaging comparing epi– and LS illumination and for determining the Strehl ratio of our system, U2OS cells were labeled with rabbit anti-lamin B1 (ab16048, Abcam) primary antibodies using a 1:1,000 dilution in 1% (w/v) BSA in PBS for two hours. Cells were then washed three times with 0.1% (v/v) Triton X-100 in PBS with a three-minute incubation period during each wash before being labeled with donkey anti-rabbit oligonucleotide-conjugated secondary antibodies (Massive Photonics, order number AB2401012) in antibody incubation buffer (Massive Photonics) for one hour. Cells were then washed three times with 1X washing buffer (Massive Photonics) in nanopure water, three times with 0.2% (v/v) Triton X-100 in PBS, and once in imaging buffer (500 mM NaCl in PBS, pH 8). 0.1 μm 580/605 nm fiducial beads (F8801, Invitrogen) at a dilution of 1:100,000 in nanopure water were then flowed in just before imaging. Finally, single-molecule data was obtained by flowing in imager strand solutions containing the complementary oligonucleotide-Cy3B dye conjugates (Massive Photonics, order number AB2401012) diluted in imaging buffer to a concentration of 0.01 nM.

For single-molecule imaging of microtubules for acquisition speed and resolution comparison (Fig 3, Supplementary Fig. 9), for determining the density breaking point of Easy-DHPSF (Supplementary Fig. 8), and for analysis method control (Supplementary Fig. 7d,e), U2OS cells were labeled with mouse anti-α-tubulin (T5168, Sigma-Aldrich) primary antibodies using a 1:500 dilution in 1% (w/v) BSA in PBS for two hours. Cells were then washed three times with 0.1% (v/v) Triton X-100 in PBS with a three-minute incubation period during each wash before being labeled with donkey anti-mouse oligonucleotide-conjugated secondary antibodies (Massive Photonics, order number AB2401012) at a dilution of 1:100 in antibody incubation buffer (Massive Photonics) for one hour. Cells were then washed three times with 1X washing buffer (Massive Photonics) in nanopure water, three times with 0.2% (v/v) Triton X-100 in PBS, and once in imaging buffer (500 mM NaCl in PBS, pH 8). Before imaging, 0.1 μm 580/605 nm fiducial beads (F8801, Invitrogen) at a dilution of 1:100,000 in nanopure water were flowed in. Finally, complimentary oligonucleotide-Cy3B dye conjugates (Massive Photonics, order number AB2401012) diluted in imaging buffer were flowed in at a concentration of 0.025 nM (standard concentration) or 0.2 nM (high concentration) just before imaging.

For resolution analysis comparing epi-versus LS illumination in the high-density regime with DECODE (Supplementary Fig. 10), U2OS cells were labeled with mouse anti-lamin A/C (sc-376248, Santa Cruz Biotechnology) primary antibodies at a dilution of 1:100 in 1% (w/v) BSA for two hours. Cells were then washed three times with 0.1% (v/v) Triton X-100 in PBS with a three-minute incubation period during each wash before being labeled with donkey anti-mouse oligonucleotide-conjugated secondary antibodies (Massive Photonics, order number AB2401012) at a dilution of 1:100 in antibody incubation buffer (Massive Photonics) for one hour. Cells were then washed three times with 1X washing buffer (Massive Photonics) in nanopure water, three times with 0.2% (v/v) Triton X-100 in PBS, and once in imaging buffer (500 mM NaCl in PBS, pH 8) before imaging. Finally, complimentary oligonucleotide-Cy3B dye conjugates (Massive Photonics, order number AB2401012) diluted in imaging buffer were flowed in at concentrations of 0.1 nM for high-density imaging.

For two-target imaging of Tomm20 and lamin A/C, before U2OS cells were seeded into the microfluidic chips, the chips were first flowed through with 3 μm carboxylated polystyrene fiducial beads (C37484, Invitrogen) diluted 1:100 in PBS and placed on a hotplate at 200°C for 1 minute and then at 100°C for 5 minutes to facilitate active stabilization (Supplementary Fig. 14). After seeding, fixation, permeabilization, and blocking, cells were labeled with rabbit anti-Tomm20 (ab186735, Abcam) and mouse anti-lamin A/C (sc-376248, Santa Cruz Biotechnology) primary antibodies at a dilution of 1:200 and 1:100, respectively, in 1% (w/v) BSA, 5% (v/v) salmon sperm ssDNA (ab229278, Abcam), and 10% donkey serum (ab7475, Abcam) in PBS for two hours. Cells were then washed three times with 0.1% (v/v) Triton X-100 in PBS with a three-minute incubation period during each wash before being labeled with donkey anti-rabbit and donkey anti-mouse oligonucleotide-conjugated secondary antibodies (Massive Photonics, order number AB2401012) at a dilution of 1:100 in antibody incubation buffer (Massive Photonics) for one hour. Cells were then washed three times with 1X washing buffer (Massive Photonics) in nanopure water, three times with 0.2% (v/v) Triton X-100 in PBS, and once in imaging buffer (500 mM NaCl in PBS, pH 8) before imaging. Before imaging, 0.2 μm 660/680 nm fiducial beads (F8807, Invitrogen) at a dilution of 1:100,000 in PBS were flowed in and allowed to settle for 5 min. Finally, complimentary oligonucleotide-Cy3B dye conjugates (Massive Photonics, order number AB2401012) diluted in imaging buffer were flowed in at concentrations of 0.04 nM for Tomm20 and 0.08 nM for lamin A/C sequentially for the two targets during imaging.

For two-target imaging of ezrin and lamin A/C, U2OS cells were labeled with rabbit anti-ezrin (ab40839, Abcam) and mouse anti-lamin A/C primary antibodies (sc-376248, Santa Cruz Biotechnology) at a dilution of 1:50 and 1:100, respectively, in 1% (w/v) BSA, 5% (v/v) salmon sperm ssDNA (ab229278, Abcam), and 10% donkey serum (ab7475, Abcam) in PBS for two hours. Cells were then washed three times with 0.1% (v/v) Triton X-100 in PBS with a three-minute incubation period during each wash before being labeled with donkey anti-rabbit and donkey anti-mouse oligonucleotide-conjugated secondary antibodies (Massive Photonics, order number AB2401012) at a dilution of 1:100 in antibody incubation buffer (Massive Photonics) for one hour. Cells were then washed three times with 1X washing buffer (Massive Photonics) in nanopure water, three times with 0.2% (v/v) Triton X-100 in PBS, and once in imaging buffer (500 mM NaCl in PBS, pH 8) before imaging. Next, 0.2 μm 660/680 nm fiducial beads at a dilution of 1:100,000 in imaging buffer were flowed in for drift correction and allowed to settle for 5 minutes. Finally, complimentary oligonucleotide-Cy3B dye conjugates (Massive Photonics, order number AB2401012) diluted in imaging buffer were flowed in at concentrations of 0.08 nM for ezrin and 0.08 nM for lamin A/C sequentially for the two targets during imaging.

For multi-target imaging of lamin B1, lamin A/C, and LAP2, U2OS cells were labeled with rabbit anti-lamin B1 (ab16048, Abcam), mouse anti-lamin A/C (sc-376248, Santa Cruz Biotechnology), and goat anti-thymopoietin (AF843, R&D Systems) primary antibodies at a dilution of 1:1000, 1:100, and 1:50, respectively, in 1% (w/v) BSA, 5% (v/v) salmon sperm ssDNA (ab229278, Abcam), and 10% donkey serum (ab7475, Abcam) in PBS for two hours. Cells were then washed three times with 0.1% (v/v) Triton X-100 in PBS with a three-minute incubation period during each wash before being labeled with donkey anti-rabbit, donkey anti-mouse, and donkey anti-goat oligonucleotide-conjugated secondary antibodies (Massive Photonics, order number AB2401012) at a dilution of 1:100 in antibody incubation buffer (Massive Photonics) for one hour. Cells were then washed three times with 1X washing buffer (Massive Photonics) in nanopure water, three times with 0.2% (v/v) Triton X-100 in PBS, and once in imaging buffer (500 mM NaCl in PBS, pH 8). Before imaging, 0.2 μm 660/680 nm fiducial beads (F8807, Invitrogen) at a dilution of 1:100,000 in PBS were flowed in. Finally, complimentary oligonucleotide-Cy3B dye conjugates (Massive Photonics, order number AB2401012) diluted in imaging buffer were flowed in at concentrations of 0.04 nM for lamin B1, 0.08 nM for lamin A/C, and 0.08 nM for LAP2 sequentially for the three targets during imaging. Multi-target sequential labeling controls are shown in Supplementary Fig. 16.

For diffraction-limited and dSTORM super-resolution imaging of stem cell aggregates, stem cells were labeled with rabbit anti-lamin B1 primary antibodies (ab16048, Abcam) using a 1:1,000 dilution in 10% (v/v) donkey serum in PBS for two hours. Cells were then washed three times with 0.1% (v/v) Triton X-100 in PBS with a three-minute incubation period during each wash before being labeled with donkey anti-rabbit secondary antibodies conjugated with dye CF568 (20098-1, Biotium) at a 1:100 dilution in 10% (v/v) donkey serum in PBS for one hour. Cells were then washed five times in 0.1% (v/v) Triton X-100 in PBS. An oxygen scavenging buffer containing 100 mM Tris-HCl, 10% (w/v) glucose, 2 μl/ml catalase, and 560 μg/ml glucose oxidase was added during imaging to reduce photobleaching, and this buffer was supplemented with 286 mM β-mercaptoethanol (BME) for dSTORM imaging.

### Imaging procedure and settings

For diffraction-limited imaging of lamin B1 in U2OS cells (Fig. 2a, Supplementary Fig. 5), cells were imaged with the 560 nm laser at ∼90 W/cm^2^ and an exposure time of 50 ms.

Single-molecule images were acquired to compare epi– and LS illumination SBR and localization precision (Fig. 2b,c, Supplementary Fig. 6), to determine the Strehl ratio of our system (Supplementary Fig. 3), to compare the speed and resolution when using Easy-DHPSF or DECODE for analysis (Fig. 3, Supplementary Figs. 8 and 9), and for epi-versus LS illumination resolution analysis in the high-density regime with DECODE (Supplementary Fig. 10). For the epi-versus LS illumination localization precision comparisons of lamin B1, the 560 nm laser was used at ∼580 W/cm^2^ for epi-illumination and at ∼800 W/cm^2^ for LS illumination using an exposure time of 100 ms to match the binding kinetics of the imager strands. 50,000 frames were acquired. To determine the Strehl ratio of our system, lamin B1 in U2OS cells was imaged with the 560 nm laser at ∼800 W/cm^2^ first at 0 µm and then at 5 µm above the coverslip. 3,000 frames at a 100 ms exposure time were acquired at each height and the resulting median photon counts were compared. For the speed and resolution comparison of microtubules, the 560 nm laser was used at ∼800 W/cm^2^, the exposure time was set to 100 ms, and 100,000 frames were acquired. For the resolution analysis of lamin A/C comparing epi-versus LS illumination with DECODE in the high emitter density regime, three cells were imaged with the 560 nm laser at ∼470 W/cm^2^ for epi-illumination and ∼450 W/cm^2^ for LS illumination at an exposure time of 100 ms. 10,000 frames were acquired for each data set and the resulting number of localizations and resolution values in the xy, xz, and yz planes were determined. After imaging, dark frames were acquired with the lasers turned off and camera shutters closed and z-scan calibration stacks of the 2-μm axial range DH-PSF in the green channel were acquired with fiducial beads (T7280, TetraSpeck, 0.2 µm, Invitrogen) spin coated in 1% (w/w) PVA (Mowiol 4-88, #17951, Polysciences Inc.) in nanopure water on a coverslip.

For two-target single-molecule imaging of mitochondria and lamin A/C (Fig. 4a,b), 50,000 frames were acquired for each target using the 560 nm laser at ∼550 W/cm^2^ and an exposure time of 100 ms. For fiducial bead drift correction, 0.2 μm 660/680 nm fiducial beads were excited using 642 nm epi-illumination at ∼3 W/cm^2^. For active stabilization with 3 μm carboxylated polystyrene fiducial beads, the 850 nm LED was used at ∼1.3 W/cm^2^ (Supplementary Fig. 14). The CMOS camera was operated at an exposure time of 10 ms with a correction speed of 400 ms for active stabilization. Imager strands for each target and washing buffer between targets were flowed through the microfluidic with a pressure-based flow control pump set to a pressure of 345 mbar, which corresponded to a calibrated flow rate of 58 µL/min. After imaging, dark frames were acquired with the lasers turned off and camera shutters closed and z-scan calibration stacks of the 12-μm axial range DH-PSF in the red channel and the 2-μm axial range DH-PSF in the green channel were acquired with fiducial beads (T7280, TetraSpeck, 0.2 µm, Invitrogen) spin coated in 1% (w/w) PVA in nanopure water on a coverslip. Registration images of both channels were acquired using the standard PSF with fiducial beads (T7280, TetraSpeck, 0.2 µm, Invitrogen) spin coated in 1% (w/w) PVA in nanopure water on a coverslip.

For two-target single-molecule imaging of ezrin and lamin A/C (Fig. 4c), 20,000 frames of ezrin and 10,000 frames of lamin A/C were acquired at each slice using the 560 nm laser at ∼ 550 W/cm^2^ and an exposure time of 100 ms. For fiducial bead drift correction, 0.2 μm 660/680 nm fiducial beads were excited using 642 nm epi-illumination at ∼3 W/cm^2^. To combat fiducial bead saturation, a 1.0 ND filter was placed in the red channel of the emission path. Imager strands for each target and washing buffer between targets were flowed through the microfluidic channel with the pressure-based flow control pump set to a pressure of 345 mbar, which corresponded to a calibrated flow rate of 58 µL/min. After imaging, dark frames were acquired with the lasers turned off and camera shutters closed and z-scan calibration stacks of the 12-μm axial range DH-PSF in the red channel and the 2-μm axial range DH-PSF in the green channel were acquired with fiducial beads (T7280, TetraSpeck, 0.2 µm, Invitrogen) spin coated in 1% (w/w) PVA in nanopure water on a coverslip. Registration images of both channels were acquired using the standard PSF with fiducial beads (T7280, TetraSpeck, 0.2 µm, Invitrogen) spin coated in 1% (w/w) PVA in nanopure water on a coverslip.

For multi-target single-molecule imaging of the three nuclear targets (Fig. 5a,b), 10,000 frames were acquired for each target at each slice using the 560 nm laser at ∼190 W/cm^2^ and an exposure time of 100 ms. For fiducial bead drift correction, 0.2 μm 660/680 nm fiducial beads were excited using 642 nm epi-illumination at ∼3 W/cm^2^. To combat fiducial bead saturation, a 1.0 ND filter was placed in the red channel of the emission path. Imager strands for each target and washing buffer between targets were flowed through the microfluidic channel with the pressure-based flow control pump set to a pressure of 345 mbar, which corresponded to a calibrated flow rate of 58 µL/min. After imaging, dark frames were acquired with the lasers turned off and camera shutters closed and z-scan calibration stacks of the 12-μm axial range DH-PSF in the red channel and the 2-μm axial range DH-PSF in the green channel were acquired with fiducial beads (T7280, TetraSpeck, 0.2 µm, Invitrogen) spin coated in 1% (w/w) PVA in nanopure water on a coverslip. Registration images of both channels were acquired using the standard PSF with fiducial beads (T7280, TetraSpeck, 0.2 µm, Invitrogen) spin coated in 1% (w/w) PVA in nanopure water on a coverslip.

For diffraction-limited imaging of stem cell aggregates comparing epi– and LS illumination (Supplementary Fig. 13), cells were imaged with the 560 nm laser at ∼ 0.2-4 W/cm^2^ for epi-illumination and ∼ 0.2-5 W/cm^2^ for LS illumination at an exposure time of 50 ms (Supplementary Fig. 13a-c). For dSTORM imaging of stem cell aggregates (Supplementary Fig. 13d-f), cells were imaged with the 560 nm laser at ∼ 2 kW/cm^2^ using LS illumination with an exposure time of 50 ms for 10,000 frames (Supplementary Fig. 13d,e) and 7,500 frames (Supplementary Fig. 13f). Blinking buffer was flowed through the microfluidic channel with the pressure-based flow control pump set to a pressure of 345 mbar, which corresponded to a calibrated flow rate of 58 µL/min.

### Data analysis

After imaging, 3D single-molecule images acquired using the DH phase mask in the green channel were imported into either Easy-DHPSF^69,70^, a MATLAB-based open-source localization software for non-overlapping emitters, or the deep learning based single-molecule detection and localization tool DECODE^12^ for overlapping emitter analysis.

2D single-molecule data for Exchange-PAINT labeling and imaging controls (Supplementary Fig. 16), for Strehl ratio determination (Supplementary Fig. 3), and for stem cell aggregate imaging with dSTORM (Supplementary Fig. 13 d-f) was analyzed in ThunderSTORM^78^, an open-source ImageJ plugin using wavelet filtering for background subtraction and a weighted least-squares fitting routine. PSF detection was performed with a local maximum fitting method with a peak intensity threshold coefficient of 2.5.

For imaging where the fiducial bead was detected in the same channel as the single-molecule data, data localized by Easy-DHPSF was drift corrected with the built in Easy-DHPSF drift correction script while DECODE-localized data was drift corrected separately with a custom MATLAB script using fiducial bead data localized in Easy-DHPSF. For two-channel whole-cell imaging, where fiducial bead data was acquired in the red channel with the 12-µm range DH-PSF and single-molecule data was acquired in the green channel with the 2-µm range DH-PSF, the 12-µm fiducial bead data was localized with a custom-edited version of Easy-DHPSF compatible with the 12-µm range DH-PSF. The 12-µm range fiducial bead data was then transformed into the green channel with a custom 2D registration code which utilized an affine transformation to map beads from one channel to another. The tracked motion of the fiducial bead was smoothed with a cubic spline fitting function and subtracted from the high-density green channel single-molecule data localized by DECODE using a custom MATLAB script. For whole-cell imaging, slices were stitched together with custom scripts which first shifted the data sets based on the position of fiducial beads that were detectable across multiple slices owing to the very long axial range of the 12-µm DH-PSF, and then corrected any residual offsets using cross-correlation between adjacent slices. Finally, localizations were filtered along a gradient from the top of each slice to avoid the appearance of harsh lines between stitched slices. An illustration of the full data analysis pipeline is shown in Supplementary Fig. 17.

To account for index mismatch between the glass coverslip and the sample, all z localizations were scaled by a factor of 0.75^55,79^ (Supplementary Fig. 18). The z compression factor of 0.75 was experimentally determined using 4 μm fluorescent beads (Fluospheres Sulfate Microspheres 4.0 μm (580/605), Invitrogen). Beads were diluted at a concentration of 1:1000 in PBS before being left to dry and adhere in an 8-well ibidi chamber (80827, Ibidi GmbH) overnight in an oven at 45°C. After adhesion, the beads were imaged in 1X PBS. Z-scans of 6 beads were performed that were each 8 μm in total height with 100 nm steps using 560 nm epi-illumination at ∼95 W/cm^2^ with a calibrated EM gain of 182 and an exposure time of 50 ms. After the z-scan was performed, an ellipse was fit to the orthogonal yz projection of each bead and the ratio between the major and minor axes of the ellipse was used to determine the experimental compression factor^80^.

Once 3D single-molecule data had been localized, drift corrected, and stitched together, it was filtered to remove weak localizations. Epi-versus LS illumination comparison data of lamin B1 (Fig. 2c and Supplementary Fig. 6) was filtered to remove any localizations with localization precisions in x, y or z greater than 100 nm, photon counts greater than 50,000, and a DH lobe separation below 4.5 or greater than 8.5 pixels. For Easy-DHPSF versus DECODE comparison data of microtubules (Fig. 3a and Supplementary Fig. 9), localizations with a localization precision greater than 20 nm in xy or greater than 30 nm in z were filtered out. The data sets were also filtered for 20-nearest neighbors with a denoise range of 0.01 to 10 to remove spurious localizations. For reconstructions and FRC resolution values without filtering of spurious localizations, please see Supplementary Figure 19. For microtubule data sets quantifying the density breaking point of Easy-DHPSF and the resulting resolution of high-density data when analyzed with DECODE compared to Easy-DHPSF (Supplementary Fig. 8), reconstructions were filtered to remove localizations with localization precisions greater than 30 nm in xy and greater than 40 nm in z. For high-density single-molecule LS and epi-illumination comparison data of lamin A/C (Supplementary Fig. 10), all data was filtered to remove localizations with localization precision greater than 30 nm. The lamin A/C and mitochondria data sets (Fig. 4a,b) were filtered to remove localizations with localization precision greater than 30 nm in xy or greater than 50 nm in z. The ezrin and lamin A/C reconstructions (Fig. 4c) were filtered to remove localizations with localization precisions greater than 50 nm in xy and greater than 100 nm in z. Whole-cell lamin B1, lamin A/C, and LAP2 data sets (Figs. 5a,b) were filtered to remove localizations with localization precision greater than 30 nm in xy and greater than 150 in z, and for 10-nearest neighbors with a denoise range of 0.01 to 10 to remove spurious localizations. Each slice was also filtered to remove localizations with z values greater than 1 µm (the upper limit of our 2-µm DH-PSF range). For reconstructions and FRC resolution values without filtering of spurious localizations, please see Supplementary Figure 19. The 2D data acquired for Strehl ratio determination was filtered to remove localizations with sigma values, the width of the Gaussian that fit the localizations, greater than 300 nm (Supplementary Fig. 3). The 2D data for the labeling and imaging controls were unfiltered (Supplementary Fig. 16). The 2D super-resolution stem cell aggregate data sets were filtered to remove localizations with photon counts greater than 5,000 (Supplementary Fig. 13 d-f).

Finally, single-molecule data was rendered using the software Vutara SRX (Bruker) for visualization, where localizations were visualized by a 3D Gaussian using point splat rendering. Microtubule reconstructions in Fig. 3a and Supplementary Fig. 19 were rendered using a 30 nm particle size diameter. Microtubule reconstructions in Supplementary Fig. 8a,c,d were rendered using a 50 nm particle size diameter. Microtubule reconstructions in Supplementary Fig. 9 were rendered using a 25 nm particle size diameter. Lamin A/C and mitochondria reconstructions in Fig. 4a,b, not including lamin A/C inset panels were rendered using a 16 nm particle size diameter. Lamin A/C inset panels in Fig. 4a were rendered using a 20 nm particle size diameter. Lamin A/C and ezrin reconstructions in Fig. 4c were rendered using a 30 nm particle size diameter. Lamin A/C, LAP2, and lamin B1 reconstructions in Fig. 5 and Supplementary Fig. 19 were rendered using a 16 nm particle size diameter. Stem cell reconstructions in Supplementary Fig. 13d were rendered using a 100 nm particle size diameter and with a 60 nm particle size diameter in Supplementary Fig. 13e,f.

For resolution analysis, the FRC was calculated in Vutara SRX and used to analyze data in the xy, yz, and xz planes (Fig. 3c,d and 5e, and Supplementary Figs. 8c,d, 9, 10, 11, and 19). A super-resolution pixel size of 8 nm was used with a threshold of 0.143 (1/7) to extract the resolution. The FRC resolution will typically improve with the number of localizations until the point it becomes limited by the localization precision and residual drift.

For DECODE training, DECODE receives experimental single-molecule data and simulates realistic single-molecule data for training. For this work, a DECODE model was trained to detect high density data acquired with the 2-µm range DH-PSF. Training was performed by feeding the DECODE environment a calibration file based on the experimental parameters of our imaging system including signal photons in the range of 0 to 18,000 photons per localization, background levels of 0 to 200 photons per pixel, a dark level of 477 A/D counts, an EM gain of 182, a conversion gain of 4.41 photoelectrons per A/D count, and the axial range of our 2-µm PSF, as well as sparse 3D single-molecule data acquired with our single-objective LS. High density data sets were then simulated based on these conditions. The DECODE model was trained on our 0.025 nM microtubule data set and converged to a Jaccard index of 0.69 after 1000 epochs. Details related to acquisition time and reconstruction time for each 3D SR reconstruction can be found in Supplementary Table 1. The DECODE model used to fit all data took ∼12 hours to train with a NVIDIA GeForce RTX 2060 12GB graphics processing unit on a Windows 10 64-bit operating system. The performance of this model was benchmarked in terms of precision by comparing the localization precision of three different fiducial beads localized in 25,241 frames and of single-molecule data from 200,000 frames using both DECODE and Easy-DHPSF (Supplementary Fig. 7). The localization precisions in x, y, and z for the three beads were extracted from Gaussian fits of the localization distributions and the localization precisions for the single-molecule data were extracted from each software. The results showed comparable values between the two approaches both for the beads and single-molecule data.

## Data availability

The single-molecule localizations generated and analyzed during the current study are available from the corresponding author upon request and from Zenodo at the following DOI: 10.5281/zenodo.12786066.

## Code availability

Calibration and fitting analysis of sparse 2-µm range DH-PSF images was performed using a modified version of the open-source Easy-DHPSF software^69,70^ (https://sourceforge.net/projects/easy-dhpsf/). Analysis of overlapping 2-µm range DH-PSF data was performed using the open-source deep learning based single-molecule localization software DECODE^12^ (https://github.com/TuragaLab/DECODE). Analysis of 2D single-molecule data was performed using the open-source ImageJ plugin ThunderSTORM^78^ (https://github.com/zitmen/thunderstorm/releases/tag/v1.3/). Active drift stabilization was achieved using an open-source active stabilization ImageJ/Micromanager plugin^13^ (https://github.com/spcoelho/Active-Stabilization/tree/master/Active%20stabilization%20ImageJ%20Plugin/ImageJ%20Plugin). The custom-written codes for red-to-green channel transformation, drift correction, slice-stitching, and the modified version of Easy-DHPSF compatible with analysis of the 12-µm range DH-PSF are available upon request.

## Acknowledgments

We thank Julia F. Love for help with the emission path construction, Armando Amador for help with initial microfluidic fabrication, Siyang Cheng and Alex Raterink for help with the active drift stabilization setup, Yuya Nakatani for help with the localization precision estimation code, and Luisa Rezende for help with stem cell culturing. This work was supported by partial financial support from the National Institute of General Medical Sciences of the National Institutes of Health grant R00GM134187 and grant R35GM155365, the Welch Foundation grant C-2064-20210327, and startup funds from the Cancer Prevention and Research Institute of Texas grant RR200025 to A.-K.G. This work was conducted in part using resources and equipment available through the Shared Equipment Authority at Rice University. We thank Dr. Timothy Gilheart, Dr. Jing Guo, and Carlos Gramajo at the Shared Equipment Authority for their input and guidance.

## Author contributions

N.S. and G.G. developed and constructed the imaging platform, performed the experiments, and analyzed the data. N.S. developed, fabricated, and characterized the microfluidic systems. G.G. cultured and labeled the cells and performed all labeling controls. A.-K.G. conceived the idea and supervised the research. All authors contributed to writing the paper.

## Corresponding author

Correspondence to Anna-Karin Gustavsson.

## Competing interests

The authors declare that they have no competing interests.

## Notes

### Competing Interest Statement

The authors have declared no competing interest.

### Summary of Updates

This version of the manuscript has been revised to add new data for statistical backing of our results, more benchmarking, and extended discussions on limitations and comparisons with other techniques. We have updated figures 1 and 4 and added new supplementary figures 3, 5, 6, 8, 9, 10, 13, 18, 19, and supplementary table 1. We have also updated supplementary figures 11, and 16 and added new text to the manuscript accordingly.

## References

1. Weiss, L. E., Love, J. F., Yoon, J., Comerci, C. J., Milenkovic, L., Kanie, T., Jackson, P. K., Stearns, T. & Gustavsson, A.-K. Chapter 4 – Single-molecule imaging in the primary cilium. in Methods Cell Biol (eds. Bravo-San Pedro, J. M. & Galluzzi, L.) vol. 176 59–83 (Academic Press, 2023).

2. Gagliano, G., Nelson, T., Saliba, N., Vargas-Hernández, S. & Gustavsson, A.-K. Light Sheet Illumination for 3D Single-Molecule Super-Resolution Imaging of Neuronal Synapses. Front. Synaptic Neurosci. 13, 761530 (2021).

3. Gustavsson, A.-K., Ghosh, R. P., Petrov, P. N., Liphardt, J. T. & Moerner, W. E. Fast and parallel nanoscale three-dimensional tracking of heterogeneous mammalian chromatin dynamics. Mol. Biol. Cell 33, 1–11 (2022).

4. Bennett, H. W., Gustavsson, A.-K., Bayas, C. A., Petrov, P. N., Mooney, N., Moerner, W. E. & Jackson, P. K. Novel fibrillar structure in the inversin compartment of primary cilia revealed by 3D single-molecule superresolution microscopy. Mol. Biol. Cell 31, 619–639 (2020).

5. Möckl, L., Pedram, K., Roy, A. R., Krishnan, V., Gustavsson, A.-K., Dorigo, O., Bertozzi, C. R. & Moerner, W. E. Quantitative super-resolution microscopy of the mammalian glycocalyx. Dev. Cell 50, 57–72 (2019).

6. Shechtman, Y., Gustavsson, A.-K., Petrov, P. N., Dultz, E., Lee, M. Y., Weis, K. & Moerner, W. E. Observation of live chromatin dynamics in cells via 3D localization microscopy using Tetrapod point spread functions. Biomed. Opt. Express 8, 5735 (2017).

7. Gustavsson, A.-K., Petrov, P. N. & Moerner, W. E. Light sheet approaches for improved precision in 3D localization-based super-resolution imaging in mammalian cells [Invited]. Opt. Express 26, 13122 (2018).

8. Jungmann, R., Avendaño, M. S., Woehrstein, J. B., Dai, M., Shih, W. M. & Yin, P. Multiplexed 3D cellular super-resolution imaging with DNA-PAINT and Exchange-PAINT. Nat. Methods 11, 313– 318 (2014).

9. van de Linde, S., Endesfelder, U., Mukherjee, A., Schüttpelz, M., Wiebusch, G., Wolter, S., Heilemann, M. & Sauer, M. Multicolor photoswitching microscopy for subdiffraction-resolution fluorescence imaging. Photochem Photobiol Sci 8, 465–469 (2009).

10. Mortensen, K. I., Churchman, L. S., Spudich, J. A. & Flyvbjerg, H. Optimized localization analysis for single-molecule tracking and super-resolution microscopy. Nat. Methods 7, 377–381 (2010).

11. Chung, K. K. H., Zhang, Z., Kidd, P., Zhang, Y., Williams, N. D., Rollins, B., Yang, Y., Lin, C., Baddeley, D. & Bewersdorf, J. Fluorogenic DNA-PAINT for faster, low-background super-resolution imaging. Nat Methods 19, 554–559 (2022).

12. Speiser, A., Müller, L.-R., Hoess, P., Matti, U., Obara, C. J., Legant, W. R., Kreshuk, A., Macke, J. H., Ries, J. & Turaga, S. C. Deep learning enables fast and dense single-molecule localization with high accuracy. Nat Methods 18, 1082–1090 (2021).

13. Coelho, S., Baek, J., Graus, M. S., Halstead, J. M., Nicovich, P. R., Feher, K., Gandhi, H., Gooding, J. J. & Gaus, K. Ultraprecise single-molecule localization microscopy enables in situ distance measurements in intact cells. Sci. Adv. 6, eaay8271 (2020).

14. Huisken, J., Swoger, J., Del Bene, F., Wittbrodt, J. & Stelzer, E. H. K. Optical sectioning deep inside live embryos by selective plane illumination microscopy. Science 305, 1007 (2004).

15. Gustavsson, A.-K., Petrov, P. N., Lee, M. Y., Shechtman, Y. & Moerner, W. E. 3D single-molecule super-resolution microscopy with a tilted light sheet. Nat. Commun. 9, 123 (2018).

16. Nelson, T., Vargas-Hernández, S., Freire, M., Cheng, S. & Gustavsson, A.-K. Multimodal illumination platform for 3D single-molecule super-resolution imaging throughout mammalian cells. Biomed. Opt. Express 15, 3050–3063 (2024).

17. Tokunaga, M., Imamoto, N. & Sakata-Sogawa, K. Highly inclined thin illumination enables clear single-molecule imaging in cells. Nat. Methods 5, 159–161 (2008).

18. Konopka, C. A. & Bednarek, S. Y. Variable-angle epifluorescence microscopy: a new way to look at protein dynamics in the plant cell cortex. Plant J. 53, 186–196 (2008).

19. Huisken, J. & Stainier, D. Y. R. Even fluorescence excitation by multidirectional selective plane illumination microscopy (mSPIM). Opt. Lett. 32, 2608 (2007).

20. Cella Zanacchi, F., Lavagnino, Z., Perrone Donnorso, M., Del Bue, A., Furia, L., Faretta, M. & Diaspro, A. Live-cell 3D super-resolution imaging in thick biological samples. Nat Methods 8, 1047– 1049 (2011).

21. Gebhardt, J. C. M., Suter, D. M., Roy, R., Zhao, Z. W., Chapman, A. R., Basu, S., Maniatis, T. & Xie, X. S. Single-molecule imaging of transcription factor binding to DNA in live mammalian cells. Nat. Methods 10, 421–426 (2013).

22. Hu, Y. S., Zhu, Q., Elkins, K., Tse, K., Li, Y., Fitzpatrick, J. A. J., Verma, I. M. & Cang, H. Light-sheet Bayesian microscopy enables deep-cell super-resolution imaging of heterochromatin in live human embryonic stem cells. Opt. Nanoscopy 2, 7 (2013).

23. Chen, B.-C., Legant, W. R., Wang, K., Shao, L., Milkie, D. E., Davidson, M. W., Janetopoulos, C., Wu, X. S., Hammer, J. A., Liu, Z., English, B. P., Mimori-Kiyosue, Y., Romero, D. P., Ritter, A. T., Lippincott-Schwartz, J., Fritz-Laylin, L., Mullins, R. D., Mitchell, D. M., Bembenek, J. N., Reymann, A.-C., Böhme, R., Grill, S. W., Wang, J. T., Seydoux, G., Tulu, U. S., Kiehart, D. P. & Betzig, E. Lattice light-sheet microscopy: Imaging molecules to embryos at high spatiotemporal resolution. Science 346, 1257998 (2014).

24. Moore, R. P., O’Shaughnessy, E. C., Shi, Y., Nogueira, A. T., Heath, K. M., Hahn, K. M. & Legant, W. R. A multi-functional microfluidic device compatible with widefield and light sheet microscopy. Lab Chip 22, 136–147 (2021).

25. Zhang, J., Zhang, M., Wang, Y., Donarski, E. & Gahlmann, A. Optically Accessible Microfluidic Flow Channels for Noninvasive High-Resolution Biofilm Imaging Using Lattice Light Sheet Microscopy. J. Phys. Chem. B 125, 12187–12196 (2021).

26. Hong, W., Wright, T., Sparks, H., Dvinskikh, L., MacLeod, K., Paterson, C. & Dunsby, C. Adaptive light-sheet fluorescence microscopy with a deformable mirror for video-rate volumetric imaging. Appl. Phys. Lett. 121, 193703 (2022).

27. Fadero, T. C., Gerbich, T. M., Rana, K., Suzuki, A., DiSalvo, M., Schaefer, K. N., Heppert, J. K., Boothby, T. C., Goldstein, B., Peifer, M., Allbritton, N. L., Gladfelter, A. S., Maddox, A. S. & Maddox, P. S. LITE microscopy: Tilted light-sheet excitation of model organisms offers high resolution and low photobleaching. J. Cell Biol. 217, 1869–1882 (2018).

28. Greiss, F., Deligiannaki, M., Jung, C., Gaul, U. & Braun, D. Single-molecule imaging in living drosophila embryos with reflected light-sheet microscopy. Biophys. J. 110, 939–946 (2016).

29. Jannasch, A., Szilagyi, S. A., Burmeister, M., Davis, Q. T., Hermsdorf, G. L., De, S. & Schäffer, E. Fast 3D imaging of giant unilamellar vesicles using reflected light-sheet microscopy with single molecule sensitivity. Journal of Microscopy 285, 40–51 (2022).

30. Legant, W. R., Shao, L., Grimm, J. B., Brown, T. A., Milkie, D. E., Avants, B. B., Lavis, L. D. & Betzig, E. High-density three-dimensional localization microscopy across large volumes. Nat. Methods 13, 359–365 (2016).

31. Tang, J. & Han, K. Y. Extended field-of-view single-molecule imaging by highly inclined swept illumination. Optica 5, 1063–1069 (2018).

32. Hung, S.-T., Cnossen, J., Fan, D., Siemons, M., Jurriens, D., Grußmayer, K., Soloviev, O., Soloviev, O., Kapitein, L. C., Smith, C. S. & Smith, C. S. SOLEIL: single-objective lens inclined light sheet localization microscopy. Biomed. Opt. Express 13, 3275–3294 (2022).

33. Dunsby, C. Optically sectioned imaging by oblique plane microscopy. Opt. Express 16, 20306–20316 (2008).

34. Sapoznik, E., Chang, B.-J., Huh, J., Ju, R. J., Azarova, E. V., Pohlkamp, T., Welf, E. S., Broadbent, D., Carisey, A. F., Stehbens, S. J., Lee, K.-M., Marín, A., Hanker, A. B., Schmidt, J. C., Arteaga, C. L., Yang, B., Kobayashi, Y., Tata, P. R., Kruithoff, R., Doubrovinski, K., Shepherd, D. P., Millett-Sikking, A., York, A. G., Dean, K. M. & Fiolka, R. P. A versatile oblique plane microscope for large-scale and high-resolution imaging of subcellular dynamics. eLife 9, e57681 (2020).

35. Sparks, H., Dent, L., Bakal, C., Behrens, A., Salbreux, G., Dunsby, C. & Dunsby, C. Dual-view oblique plane microscopy (dOPM). *Biomed. Opt. Express*, BOE 11, 7204–7220 (2020).

36. Chen, B., Chang, B.-J., Roudot, P., Zhou, F., Sapoznik, E., Marlar-Pavey, M., Hayes, J. B., Brown, P. T., Zeng, C.-W., Lambert, T., Friedman, J. R., Zhang, C.-L., Burnette, D. T., Shepherd, D. P., Dean, K. M. & Fiolka, R. P. Resolution doubling in light-sheet microscopy via oblique plane structured illumination. Nat Methods 19, 1419–1426 (2022).

37. Chen, B., Chang, B.-J., Zhou, F. Y., Daetwyler, S., Sapoznik, E., Nanes, B. A., Terrazas, I., Gihana, G. M., Castro, L. P., Chan, I. S., Conacci-Sorrell, M., Dean, K. M., Millett-Sikking, A., York, A. G. & Fiolka, R. Increasing the field-of-view in oblique plane microscopy via optical tiling. *Biomed. Opt. Express*, BOE 13, 5616–5627 (2022).

38. Moore, R. P., O’Shaughnessy, E. C., Shi, Y., Nogueira, A. T., Heath, K. M., Hahn, K. M. & Legant, W. R. A multi-functional microfluidic device compatible with widefield and light sheet microscopy. Lab Chip 22, 136–147 (2022).

39. Gomez-Cruz, C., Laguna, S., Bachiller-Pulido, A., Quilez, C., Cañadas-Ortega, M., Albert-Smet, I., Ripoll, J. & Muñoz-Barrutia, A. Single Plane Illumination Microscopy for Microfluidic Device Imaging. Biosensors 12, 1110 (2022).

40. Hedde, P. N., Le, B. T., Gomez, E. L., Duong, L., Steele, R. E. & Ahrar, S. SPIM-Flow: An Integrated Light Sheet and Microfluidics Platform for Hydrodynamic Studies of Hydra. Biology 12, 116 (2023).

41. Zhu, T., Nie, J., Yu, T., Zhu, D., Huang, Y., Chen, Z., Gu, Z., Tang, J., Li, D. & Fei, P. Large-scale high-throughput 3D culture, imaging, and analysis of cell spheroids using microchip-enhanced light-sheet microscopy. Biomed. Opt. Express 14, 1659–1669 (2023).

42. Galland, R., Grenci, G., Aravind, A., Viasnoff, V., Studer, V. & Sibarita, J.-B. 3D high– and super-resolution imaging using single-objective SPIM. Nat. Methods 12, 641–644 (2015).

43. Ponjavic, A., Ye, Y., Laue, E., Lee, S. F. & Klenerman, D. Sensitive light-sheet microscopy in multiwell plates using an AFM cantilever. Biomed. Opt. Express 9, 5863–5880 (2018).

44. Beghin, A., Grenci, G., Sahni, G., Guo, S., Rajendiran, H., Delaire, T., Mohamad Raffi, S. B., Blanc, D., de Mets, R., Ong, H. T., Galindo, X., Monet, A., Acharya, V., Racine, V., Levet, F., Galland, R., Sibarita, J.-B. & Viasnoff, V. Automated high-speed 3D imaging of organoid cultures with multi-scale phenotypic quantification. Nat Methods 19, 881–892 (2022).

45. Rames, M. J., Kenison, J. P., Heineck, D., Civitci, F., Szczepaniak, M., Zheng, T., Shangguan, J., Zhang, Y., Tao, K., Esener, S. & Nan, X. Multiplexed and Millimeter-Scale Fluorescence Nanoscopy of Cells and Tissue Sections via Prism-Illumination and Microfluidics-Enhanced DNA-PAINT. Chemical & Biomedical Imaging 1, 817–830 (2023).

46. Meddens, M. B. M., Liu, S., Finnegan, P. S., Edwards, T. L., James, C. D. & Lidke, K. A. Single objective light-sheet microscopy for high-speed whole-cell 3D super-resolution. Biomed. Opt. Express 7, 2219–2236 (2016).

47. Galgani, T., Fedala, Y., Zapata, R., Caccianini, L., Viasnoff, V., Sibarita, J.-B., Galland, R., Dahan, M. & Hajj, B. Selective volumetric excitation and imaging for single molecule localization microscopy in multicellular systems. 2022.12.02.518828 Preprint at 10.1101/2022.12.02.518828 (2022).

48. Gustavsson, A.-K., Adiels, C. B., Mehlig, B. & Goksör, M. Entrainment of heterogeneous glycolytic oscillations in single cells. Sci. Rep. 5, 1–7 (2015).

49. Gustavsson, A.-K., van Niekerk, D. D., Adiels, C. B., Kooi, B., Goksör, M. & Snoep, J. L. Allosteric regulation of phosphofructokinase controls the emergence of glycolytic oscillations in isolated yeast cells. FEBS J. 281, 2784–2793 (2014).

50. Gustavsson, A.-K., van Niekerk, D. D., Adiels, C. B., du Preez, F. B., Goksör, M. & Snoep, J. L. Sustained glycolytic oscillations in individual isolated yeast cells. FEBS J. 279, 2837–2847 (2012).

51. van Niekerk, D. D., Gustavsson, A.-K., Mojica-Benavides, M., Adiels, C. B., Goksör, M. & Snoep, J. L. Phosphofructokinase controls the acetaldehyde-induced phase shift in isolated yeast glycolytic oscillators. Biochem. J. 476, 353–363 (2019).

52. Gustavsson, A.-K. Heterogeneity of glycolytic oscillatory behaviour in individual yeast cells. FEBS Lett. 588, 3–7 (2014).

53. Gustavsson, A., Banaeiyan, A. A., Niekerk, D. D., Snoep, J. L., Adiels, C. B. & Goksör, M. Studying Glycolytic Oscillations in Individual Yeast Cells by Combining Fluorescence Microscopy with Microfluidics and Optical Tweezers. Curr. Prot. Cell Biol. 82, 1–26 (2019).

54. Kao, H. P. & Verkman, A. S. Tracking of single fluorescent particles in three dimensions: use of cylindrical optics to encode particle position. Biophys. J. 67, 1291–1300 (1994).

55. Huang, B., Wang, W., Bates, M. & Zhuang, X. Three-dimensional super-resolution imaging by stochastic optical reconstruction microscopy. Science 319, 810–813 (2008).

56. Lew, M. D., Lee, S. F., Badieirostami, M. & Moerner, W. E. Corkscrew point spread function for far-field three-dimensional nanoscale localization of pointlike objects. Opt. Lett. 36, 202 (2011).

57. Backer, A. S., Backlund, M. P., von Diezmann, A. R., Sahl, S. J. & Moerner, W. E. A bisected pupil for studying single-molecule orientational dynamics and its application to three-dimensional super-resolution microscopy. Appl. Phys. Lett. 104, 193701 (2014).

58. Jia, S., Vaughan, J. C. & Zhuang, X. Isotropic three-dimensional super-resolution imaging with a self-bending point spread function. Nat. Photonics 8, 302–306 (2014).

59. Shechtman, Y., Sahl, S. J., Backer, A. S. & Moerner, W. E. Optimal Point Spread Function Design for 3D Imaging. Phys. Rev. Lett. 113, 133902 (2014).

60. Shechtman, Y., Weiss, L. E., Backer, A. S., Sahl, S. J. & Moerner, W. E. Precise three-dimensional scan-free multiple-particle tracking over large axial ranges with tetrapod point spread functions. Nano Lett. 15, 4194–4199 (2015).

61. Pavani, S. R. P., Thompson, M. A., Biteen, J. S., Lord, S. J., Liu, N., Twieg, R. J., Piestun, R. & Moerner, W. E. Three-dimensional, single-molecule fluorescence imaging beyond the diffraction limit by using a double-helix point spread function. Proc. Natl. Acad. Sci. USA 106, 2995–2999 (2009).

62. Nakatani, Y., Gaumer, S., Shechtman, Y. & Gustavsson, A.-K. Long axial-range double-helix point spread functions for 3D volumetric super-resolution imaging. bioRxiv 2024.07.31.605907 (2024) doi:10.1101/2024.07.31.605907.

63. Hajj, B., Wisniewski, J., El Beheiry, M., Chen, J., Revyakin, A., Wu, C. & Dahan, M. Whole-cell, multicolor superresolution imaging using volumetric multifocus microscopy. Proceedings of the National Academy of Sciences 111, 17480–17485 (2014).

64. Zhang, P., Liu, S., Chaurasia, A., Ma, D., Mlodzianoski, M. J., Culurciello, E. & Huang, F. Analyzing complex single-molecule emission patterns with deep learning. Nat Methods 15, 913–916 (2018).

65. Zelger, P., Kaser, K., Rossboth, B., Velas, L., Schütz, G. J. & Jesacher, A. Three-dimensional localization microscopy using deep learning. Opt. Express 26, 33166 (2018).

66. Nehme, E., Weiss, L. E., Michaeli, T. & Shechtman, Y. Deep-STORM: super-resolution single-molecule microscopy by deep learning. Optica 5, 458 (2018).

67. Narayanasamy, K. K., Rahm, J. V., Tourani, S. & Heilemann, M. Fast DNA-PAINT imaging using a deep neural network. Nat Commun 13, 5047 (2022).

68. Heilemann, M., van de Linde, S., Schüttpelz, M., Kasper, R., Seefeldt, B., Mukherjee, A., Tinnefeld, P. & Sauer, M. Subdiffraction-resolution fluorescence imaging with conventional fluorescent probes. Angew. Chem. Int. Ed. 47, 6172–6176 (2008).

69. Lew, M., von Diezmann*, A. R. S. & Moerner, W. E. Easy-DHPSF open-source software for three-dimensional localization of single molecules with precision beyond the optical diffraction limit. Protocol Exchange (2013) doi:10.1038/protex.2013.026.

70. Bayas, C. A., Diezmann, A. von, Gustavsson, A.-K. & Moerner, W. E. Easy-DHPSF 2.0: open-source software for three-dimensional localization and two-color registration of single molecules with nanoscale accuracy. Prot. Exch. (2019) doi:10.21203/rs.2.9151/v2.

71. Eklund, A. S., Ganji, M., Gavins, G., Seitz, O. & Jungmann, R. Peptide-PAINT Super-Resolution Imaging Using Transient Coiled Coil Interactions. Nano Lett. 20, 6732–6737 (2020).

72. Oi, C., Gidden, Z., Holyoake, L., Kantelberg, O., Mochrie, S., Horrocks, M. H. & Regan, L. LIVE-PAINT allows super-resolution microscopy inside living cells using reversible peptide-protein interactions. *Commun*. Biol. 3, 458 (2020).

73. Lukinavičius, G., Umezawa, K., Olivier, N., Honigmann, A., Yang, G., Plass, T., Mueller, V., Reymond, L., Corrêa Jr, I. R., Luo, Z.-G., Schultz, C., Lemke, E. A., Heppenstall, P., Eggeling, C., Manley, S. & Johnsson, K. A near-infrared fluorophore for live-cell super-resolution microscopy of cellular proteins. Nat. Chem. 5, 132–139 (2013).

74. Lukinavičius, G., Reymond, L., D’Este, E., Masharina, A., Göttfert, F., Ta, H., Güther, A., Fournier, M., Rizzo, S., Waldmann, H., Blaukopf, C., Sommer, C., Gerlich, D. W., Arndt, H.-D., Hell, S. W. & Johnsson, K. Fluorogenic probes for live-cell imaging of the cytoskeleton. Nat. Methods 11, 731–733 (2014).

75. Butkevich, A. N., Mitronova, G. Yu., Sidenstein, S. C., Klocke, J. L., Kamin, D., Meineke, D. N. H., D’Este, E., Kraemer, P.-T., Danzl, J. G., Belov, V. N. & Hell, S. W. Fluorescent Rhodamines and Fluorogenic Carbopyronines for Super-Resolution STED Microscopy in Living Cells. Angew. Chem. Int. Ed. 55, 3290–3294 (2016).

76. Kessler, L. F., Balakrishnan, A., Deußner-Helfmann, N. S., Li, Y., Mantel, M., Glogger, M., Barth, H.-D., Dietz, M. S. & Heilemann, M. Self-quenched Fluorophore Dimers for DNA-PAINT and STED Microscopy. Angew. Chem. Int. Ed. 62, e202307538 (2023).

77. Halpern, A. R., Howard, M. D. & Vaughan, J. C. Point by Point: An Introductory Guide to Sample Preparation for Single-Molecule, Super-Resolution Fluorescence Microscopy: Sample Preparation for Single-Molecule Super-Resolution Fluorescence Microscopy. Curr. Prot. Chem. Biol. 7, 103–120 (2015).

78. Ovesný, M., Křížek, P., Borkovec, J., Švindrych, Z. & Hagen, G. M. ThunderSTORM: a comprehensive ImageJ plug-in for PALM and STORM data analysis and super-resolution imaging. Bioinformatics 30, 2389–2390 (2014).

79. Li, Y., Mund, M., Hoess, P., Deschamps, J., Matti, U., Nijmeijer, B., Sabinina, V. J., Ellenberg, J., Schoen, I. & Ries, J. Real-time 3D single-molecule localization using experimental point spread functions. Nat Methods 15, 367–369 (2018).

80. Bratton, B. P. & Shaevitz, J. W. Simple Experimental Methods for Determining the Apparent Focal Shift in a Microscope System. PLoS ONE 10, e0134616 (2015).

