## Supplementary Information for "Whole-cell multi-target single-molecule super-resolution imaging in 3D with microfluidics and a single-objective tilted light sheet"

### Supplementary Figures

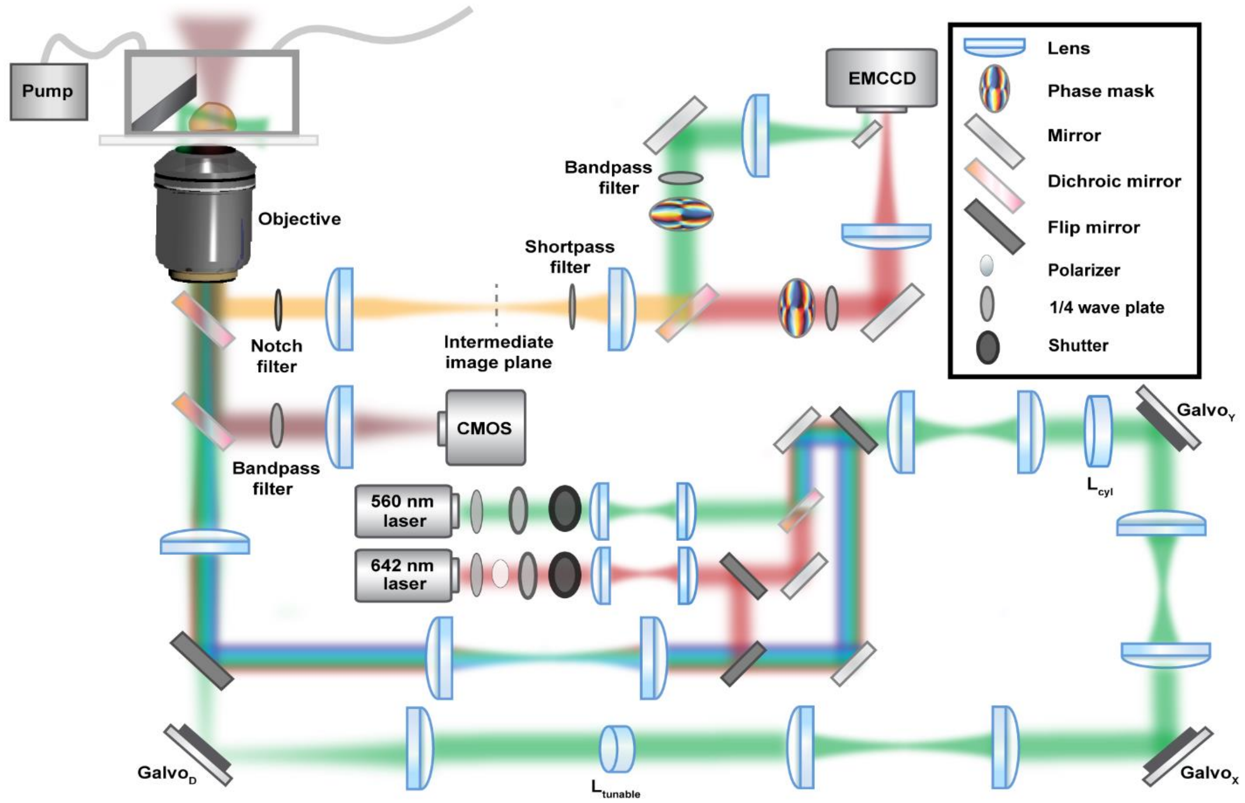


**Supplementary Fig. 1. Schematic of the optical setup.** Epi- and light sheet (LS) illumination paths are separated by a flip mirror. In the LS illumination path, the laser beam is directed to a cylindrical lens, $L_{cyl}$, where the beam is focused in one dimension to form the LS. The beam is then reflected by a galvanometric mirror, $Galvo_Y$, that is conjugated to the back focal plane of the objective and steers the LS in the y direction, and a galvanometric mirror, $Galvo_X$, that is conjugated to the back focal plane of the objective and steers the LS in the x direction. The focus of the beam is tuned using the tunable lens, $L_{tunable}$, conjugated to the back focal plane of the objective. The LS is dithered with a galvanometric mirror, $Galvo_D$, conjugated to the sample plane. Emitted light is collected through a two-channel 4*f* system with a transmissive dielectric phase mask and imaged on an EMCCD camera. Transmission light is used to image polystyrene beads on a CMOS camera for active stabilization. The schematic is not drawn to scale.

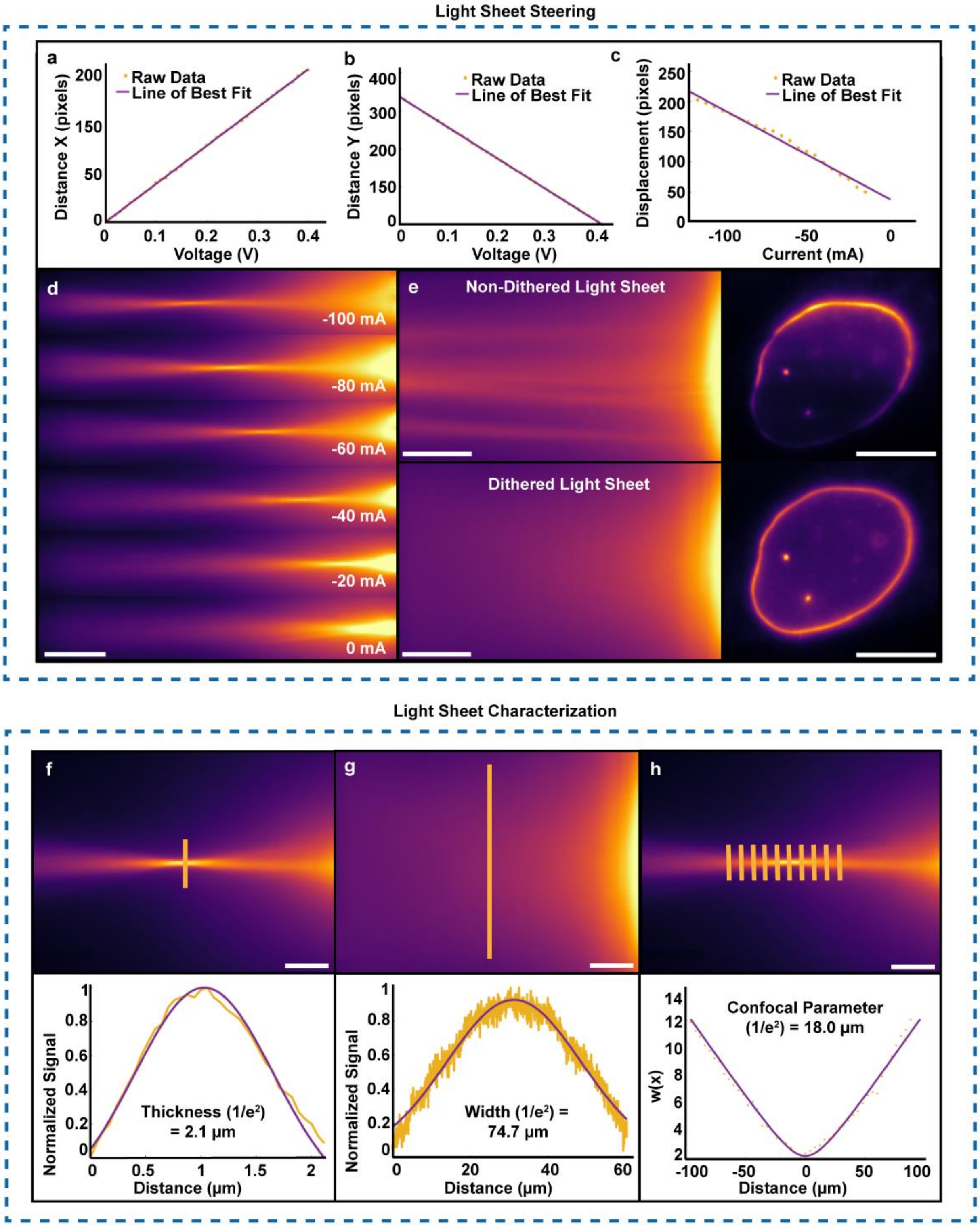


**Supplementary Fig. 2. Light sheet (LS) steering and characterization. a** Graph showing the translation in x as a function of voltage for the galvanometric X mirror. The galvanometric mirror translated the LS in a linear fashion in the x direction, where 0.01 V corresponded to a LS translation of approximately 3.86 pixels, or 0.61 μm (where 1 pixel = 0.159 μm). **b** Graph showing the LS translation in y as a function of voltage for the galvanometric Y mirror. The galvanometric mirror translated the LS in a linear fashion in the y direction, where 0.01 V corresponded to a LS translation of approximately 8.02 pixels, or 1.28 µm (where 1 pixel = 0.159 µm). **c** Graph and **d** corresponding images showing the LS focus displacement as a function of current to the tunable lens when imaged in fluorescent solution. The tunable lens was modulated with currents from 0 mA to -100 mA in increments of 20 mA, where 1 mA corresponded to moving the focus of the LS approximately 0.23 µm. Scale bar 10 µm. **e** LS illumination without and with dithering. The top row shows striping effects from a non-dithered LS as seen in fluorescent solution (left) and when imaging a U2OS cell labeled for lamin B1 (right). The bottom row demonstrates the homogenous illumination achieved by dithering the LS at a half-angle of 20° in the LS plane and a frequency of 100 Hz as seen in fluorescent solution (left) and when imaging the same cell labeled for lamin B1 (right). Scale bars 10 µm. **f** The thin end of the LS imaged in a fluorescent solution and corresponding line scan (yellow) and Gaussian fit (purple) are shown in the graph, demonstrating a thickness ($1/e^2$ diameter) of the LS of 2.1 µm, corresponding to a beam waist radius of ~1.1 µm. Scale bar 10 µm. **g** The width of the LS imaged in a fluorescent solution and corresponding line scan (yellow) and Gaussian fit (purple) are shown in the graph, demonstrating a width ($1/e^2$ full width) of the LS of 74.7 µm. Scale bar 10 µm. **h** The thin end of the LS imaged in a fluorescent solution and line scans (yellow) showing examples of where the LS thicknesses were extracted (yellow data points in the graph) along the optical axis, together with a fit (purple) to extract the confocal parameter. This fit resulted in a value of the confocal parameter ($1/e^2$) of 18.0 µm. Scale bar 10 µm.

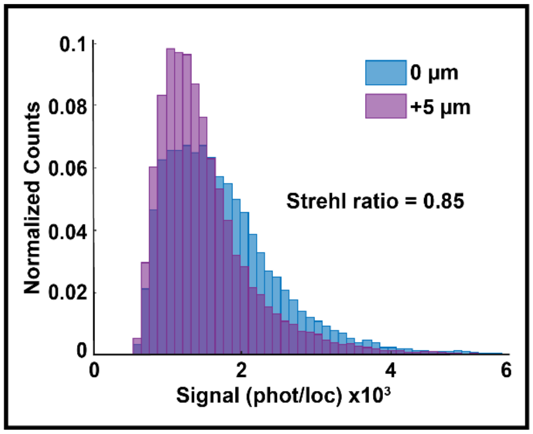


**Supplementary Fig. 3. Strehl ratio experimental determination.** Single-molecule localization signal histograms from 3,000 frames of single-molecule data of nuclear lamina protein lamin B1 acquired at 0 µm and 5 µm above the coverslip. By comparing the median localization signal at 5 µm (1,355 photons per localization) to the median localization signal at 0 µm (1,603 photons per localization), the Strehl ratio was determined to be 0.85.

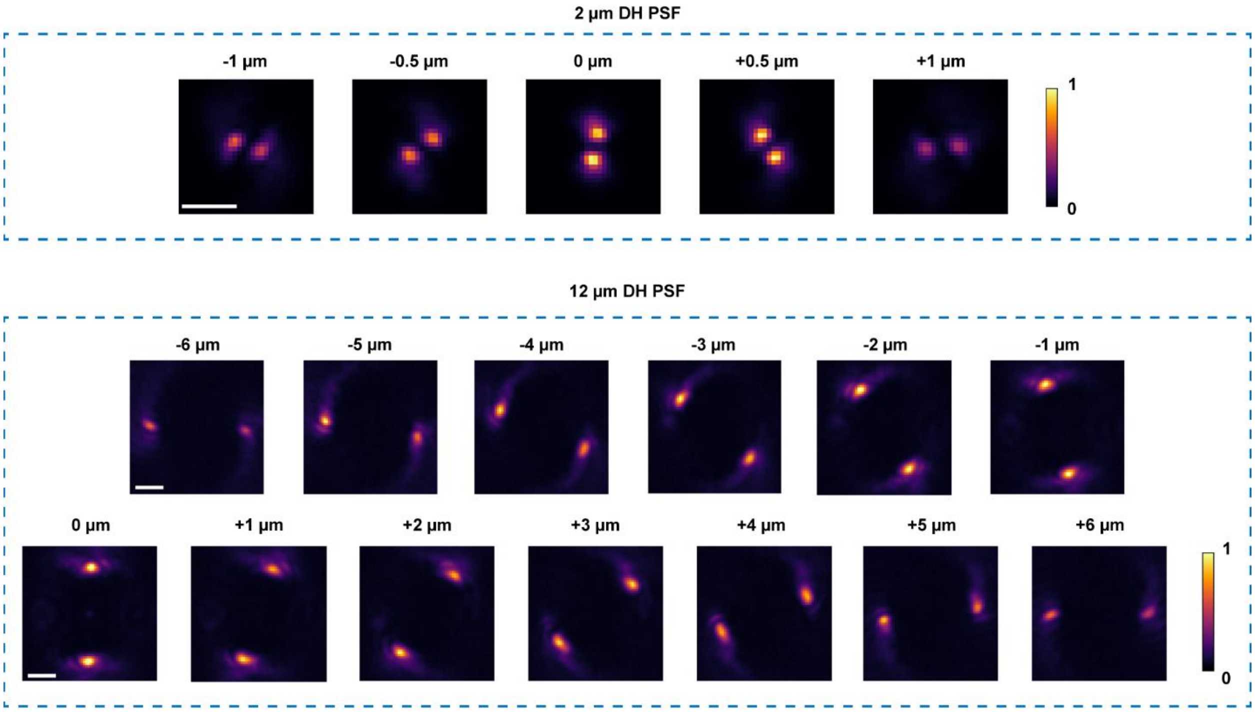


**Supplementary Fig. 4. Experimental images of the double-helix point spread functions (DH-PSFs).** The experimental short-range DH-PSF has an axial range of ~2 μm and the long-range DH-PSF has an axial range of ~12 μm. The PSFs were implemented using a transmissive dielectric phase mask in the green emission channel for the short-range DH-PSF and in the red emission channel for the long-range DH-PSF. The images show the PSF of a fluorescent bead on a coverslip imaged using the different phase masks in the corresponding channels while scanning axially using a piezo stage. Scale bars are 2 μm for the short-range DH-PSF and 3 μm for the long-range DH-PSF. The colorbars show intensity normalized independently for each image.

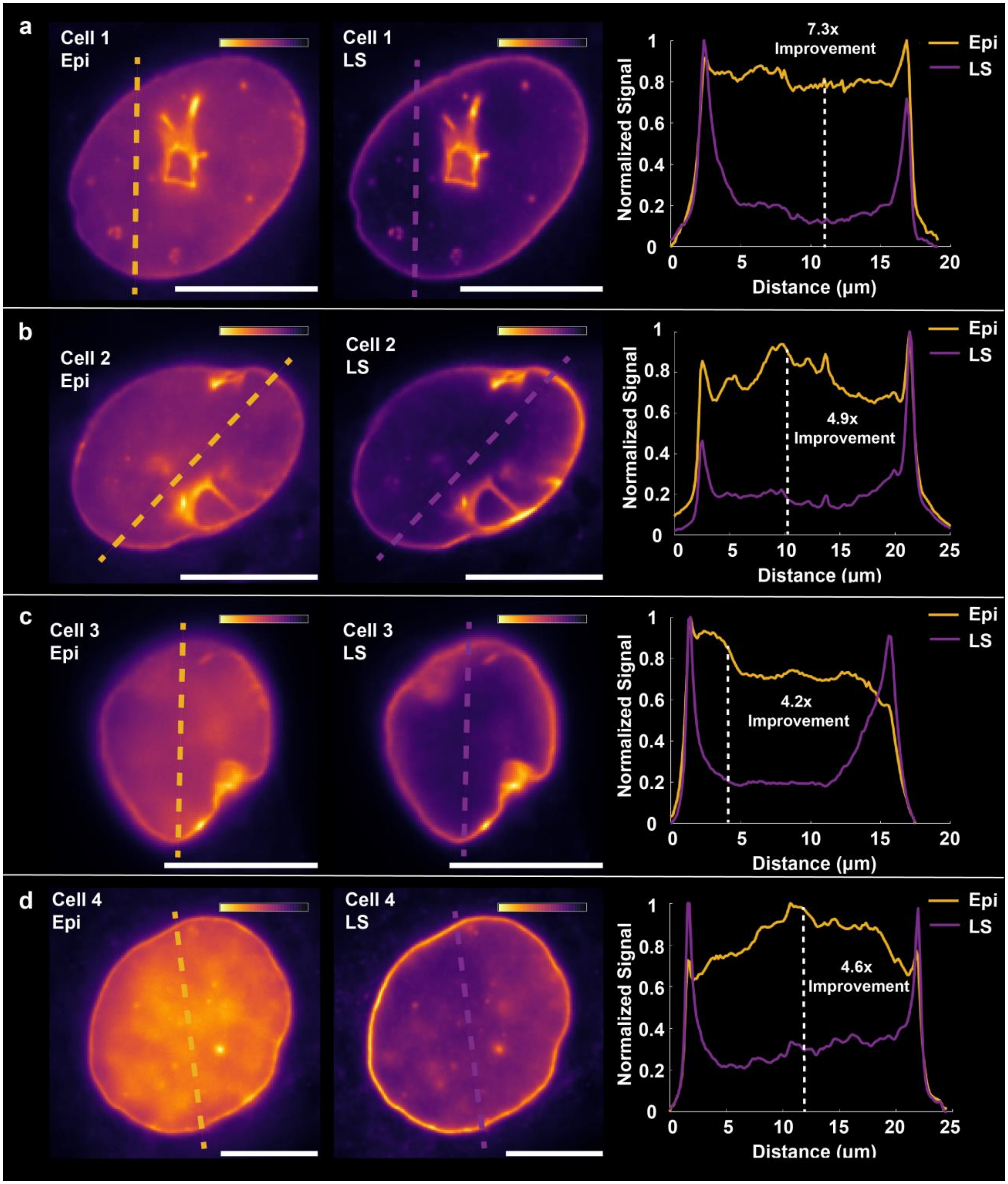


**Supplementary Fig. 5. Diffraction-limited images of lamin B1 excited with epi- or light sheet (LS) illumination.** Graphs show line scans demonstrating the consistent signal-to-background ratio (SBR) improvement with LS compared to epi-illumination. SBR improvement is variable between samples, ranging from >4x to >7x. Scale bars 10 µm. The colorbars show intensity normalized independently for each image.

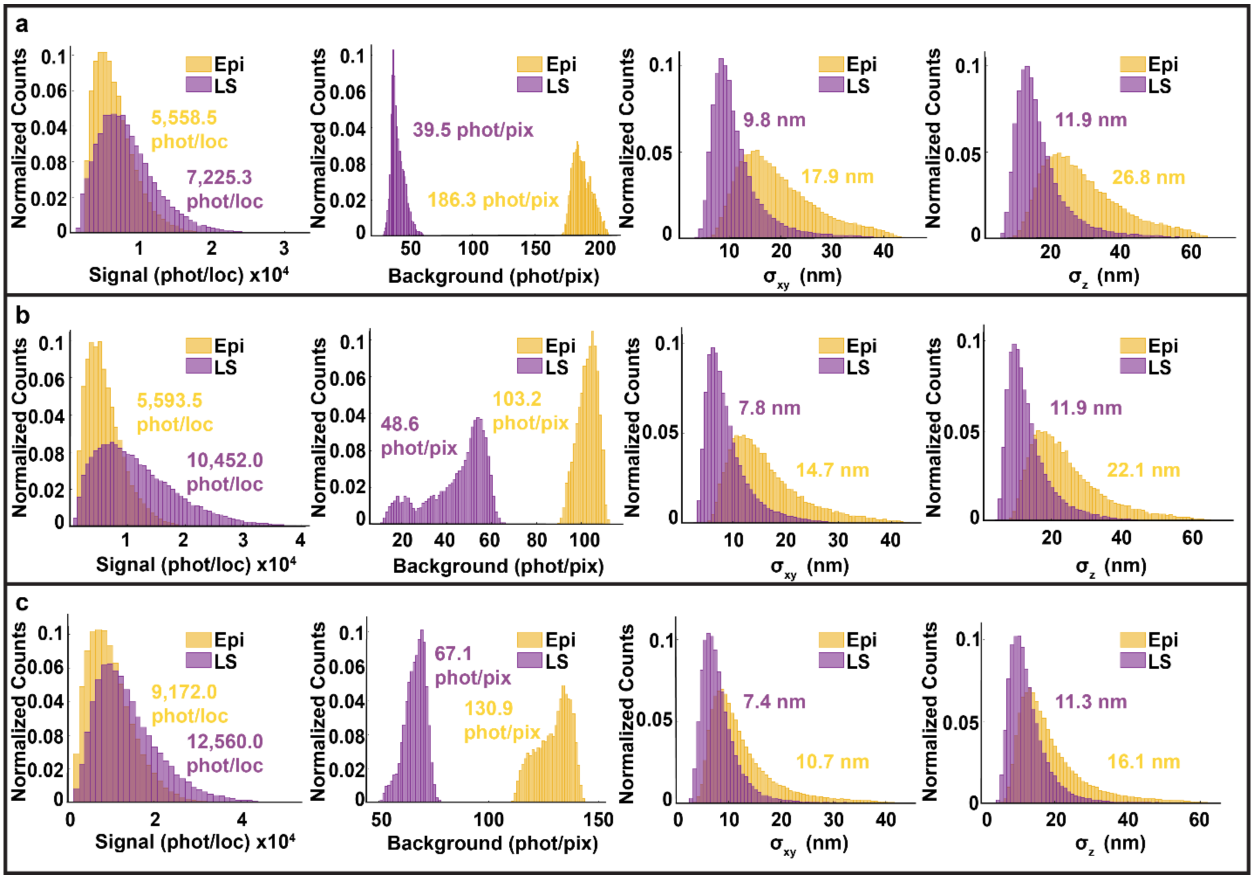


**Supplementary Fig. 6. Three technical replicates of the 3D single-molecule super-resolution imaging of lamin B1 shown in Figure 2c. a-c** Epi-illumination and light sheet (LS) illumination imaging over 10,000 frames of three different cells labeled for lamin B1 demonstrating consistent improvements in the localization precision due to background reduction when using LS illumination (purple) compared to epi-illumination (yellow). The median values of the distributions are indicated for each histogram.

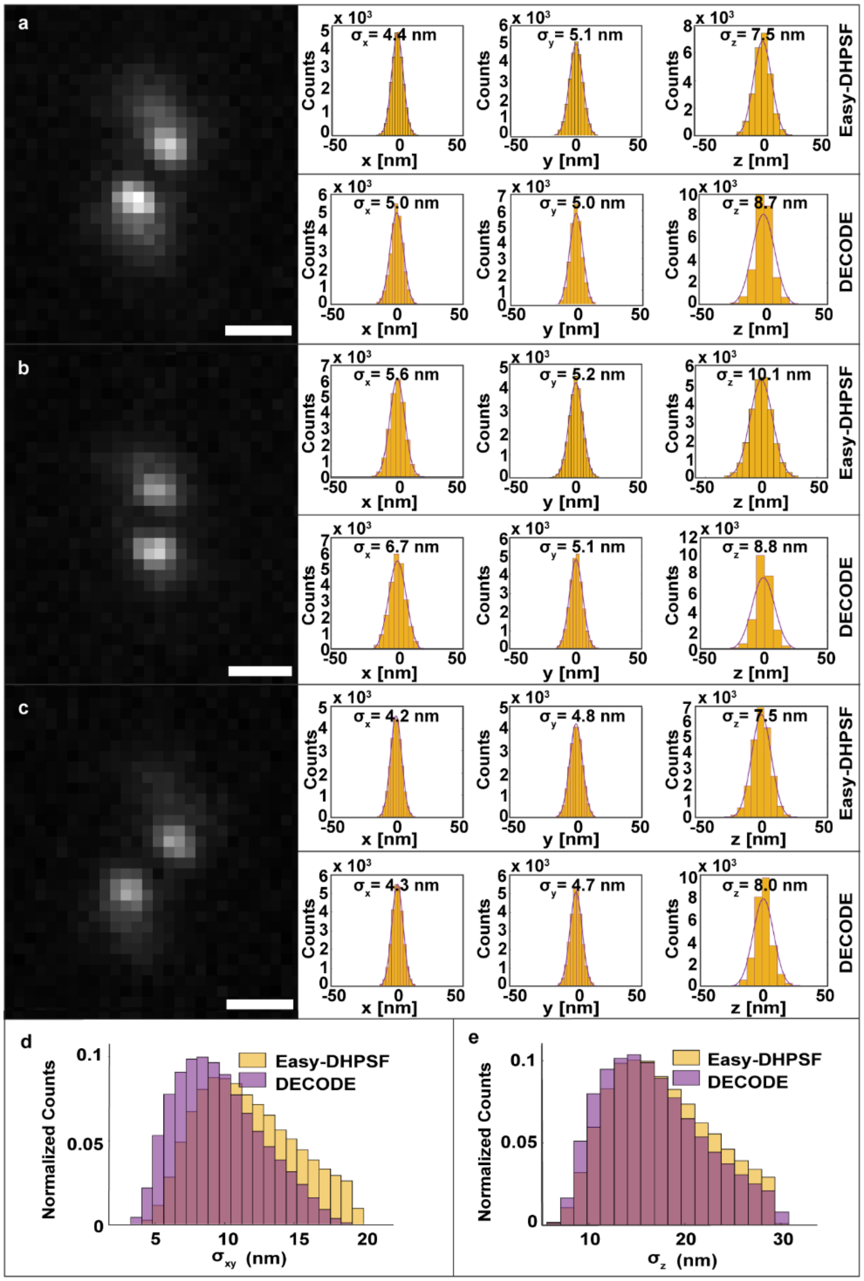


**Supplementary Fig. 7. Easy-DHPSF and DECODE localization precision analysis of fluorescent beads and single-molecule data. a-c** Three different fiducial beads imaged using the 2-µm double-helix PSF were localized for 25,241 frames by either Easy-DHPSF (top row) or DECODE (bottom row). Their bead localization distributions were plotted in the x, y and z directions, respectively, to determine the localization precision of data analyzed with each software. Localization precision values (σ) extracted from Gaussian fits of the localization distributions are shown above each histogram. Scale bars 1 µm. **d**,**e** Easy-DHPSF and DECODE localization precision analysis of 3D single-molecule data. Easy-DHPSF and DECODE were used to analyze the same microtubule single-molecule data set acquired using the 2-µm double-helix PSF, resulting in comparable **d** lateral and **e** axial localization precision values. The median localization precision values in xy/z were found to be 11.2/17.0 nm for Easy-DHPSF and 9.3/16.0 nm for DECODE after 200,000 frames. Both data sets were filtered for axial localization precision less than 20 nm and lateral localization precision less than 30 nm.

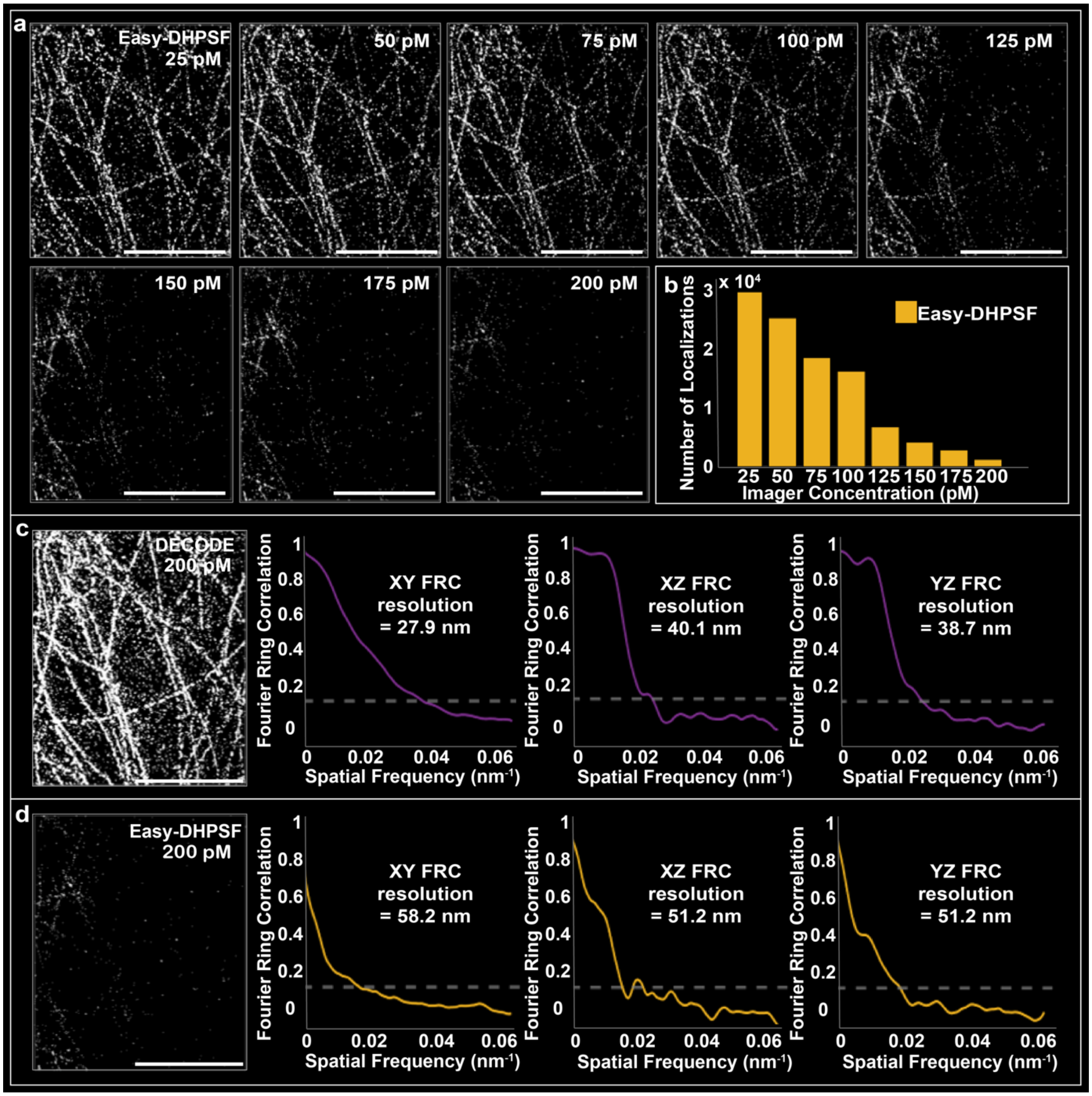


**Supplementary Fig. 8. Density breaking point for 3D single-molecule super-resolution imaging analysis with Easy-DHPSF, and comparison of the number of localizations and the resolution at overlapping-emitter densities when analyzing with Easy-DHPSF and DECODE. a** 3D single-molecule super-resolution reconstructions of microtubules imaged with increasing imager strand concentrations and analyzed with Easy-DHPSF, simulated by summing frames from a sparse single-molecule data set, demonstrating the loss of localizations as the emitter density increases. A total of 25,000 frames after summation were analyzed for each concentration. Scale bars 5 µm. **b** Quantification of the number of localizations from the images in a, demonstrating a steady decrease in detected emitters after 25 pM when analyzed with Easy-DHPSF. **c** Reconstruction of the 200 pM imager strand concentration microtubule data set from a, when analyzed with DECODE, and resulting Fourier ring correlation (FRC) resolution analysis in the xy, xz, and yz planes, respectively. **d** Reconstruction of the 200 pM imager strand concentration microtubule data set from a, analyzed with Easy-DHPSF, and resulting FRC resolution analysis in the xy, xz, and yz planes, respectively. For the same data set, DECODE detected 108,254 localizations, while EasyDHPSF detected 1,222 localizations. The FRC curves demonstrate improved resolution when analyzing with DECODE compared to Easy-DHPSF. Scale bars 5 µm.

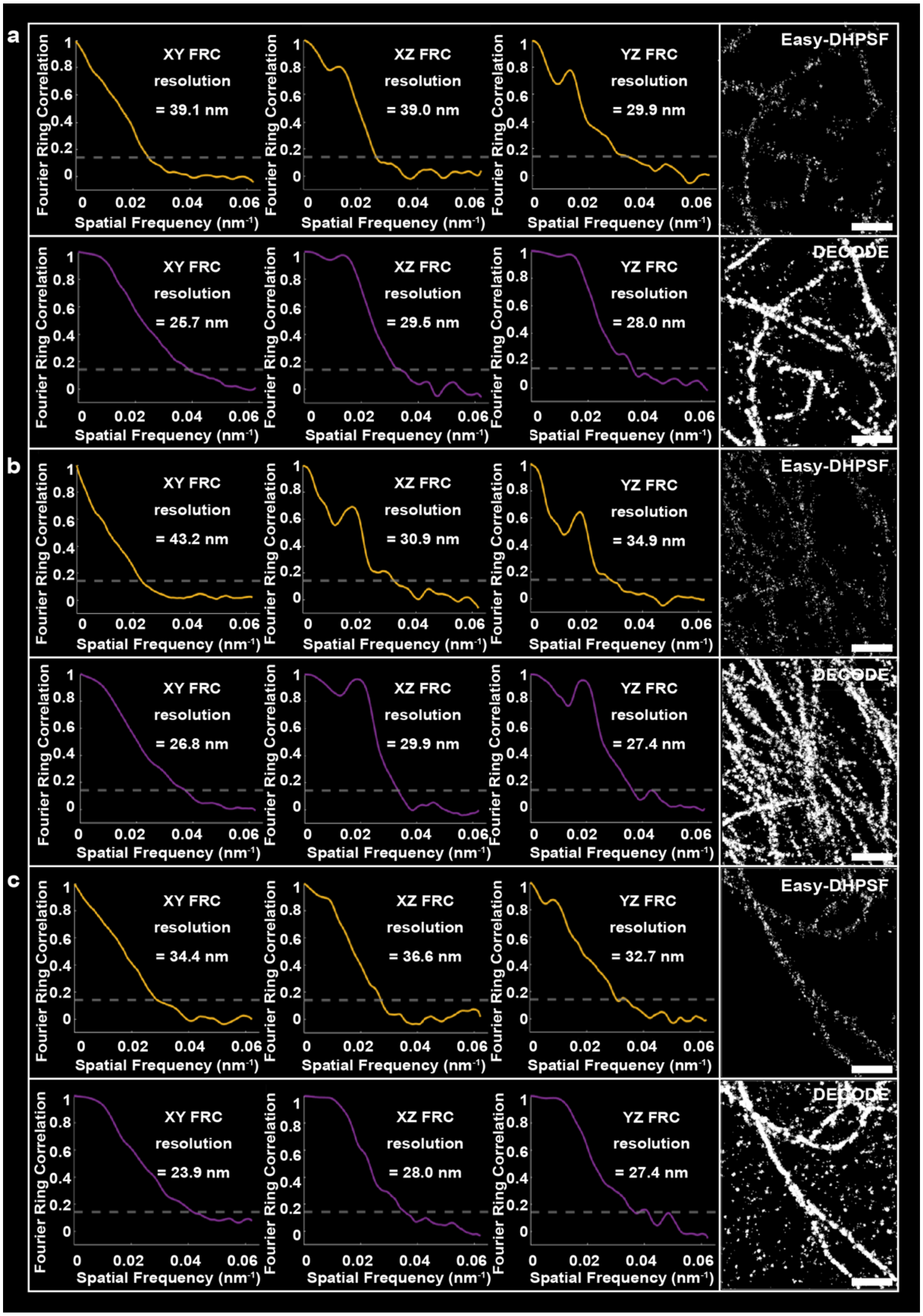


**Supplementary Fig. 9. Three technical replicates of the 3D single-molecule super-resolution imaging of microtubules shown in Figure 3c,d. a-c** Consistent resolution improvement shown by Fourier ring correlation (FRC) analysis in the xy, xz, and yz planes, respectively, for three samples of 3D single-molecule super-resolution light sheet imaging of microtubules imaged at an imager strand concentration of 0.025 nM and analyzed with Easy-DHPSF (yellow) and an imager strand concentration of 0.2 nM and analyzed with DECODE (purple). Scale bars 1 µm.

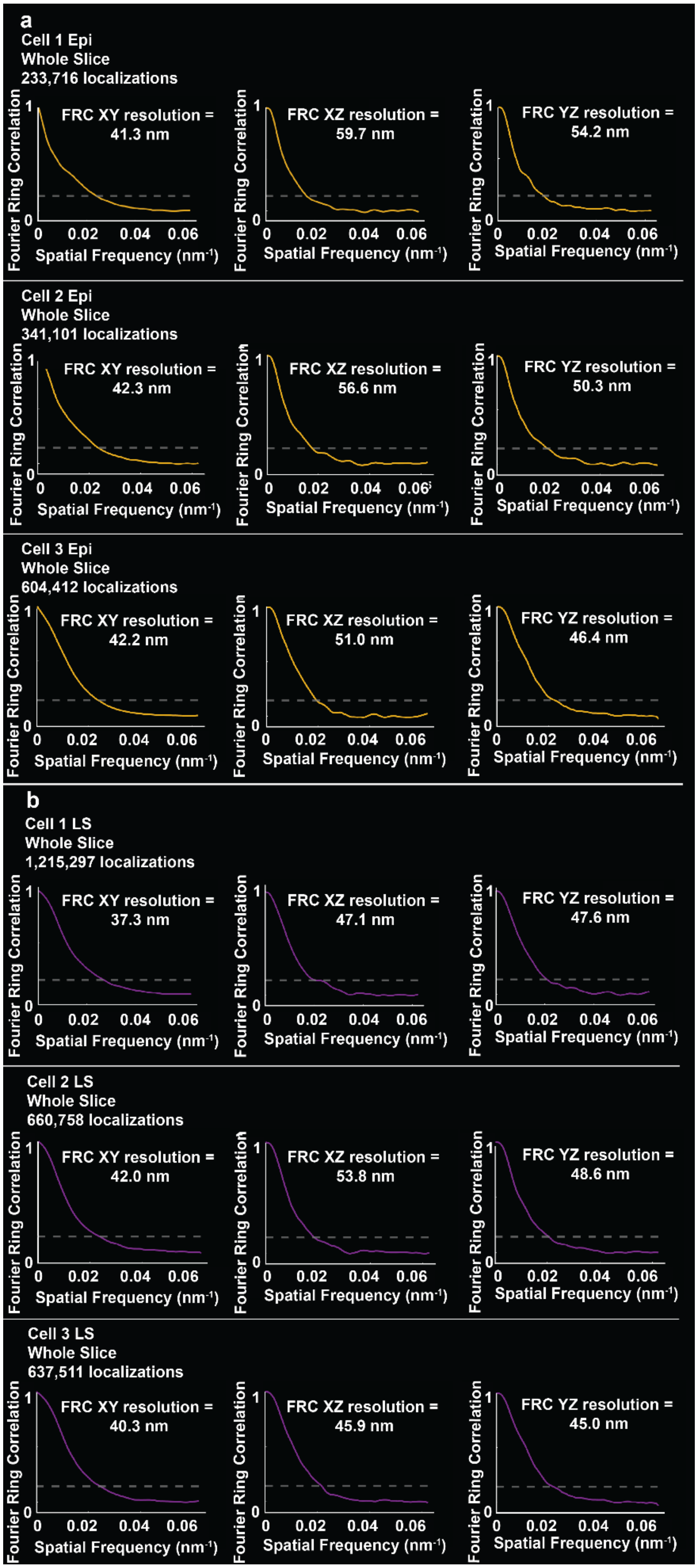
a
Cell 1 Epi
Whole Slice
233,716 localizations
FRC XY resolution = 41.3 nm
FRC XZ resolution = 59.7 nm
FRC YZ resolution = 54.2 nm
Cell 2 Epi
Whole Slice
341,101 localizations
FRC XY resolution = 42.3 nm
FRC XZ resolution = 56.6 nm
FRC YZ resolution = 50.3 nm
Cell 3 Epi
Whole Slice
604,412 localizations
FRC XY resolution = 42.2 nm
FRC XZ resolution = 51.0 nm
FRC YZ resolution = 46.4 nm
b
Cell 1 LS
Whole Slice
1,215,297 localizations
FRC XY resolution = 37.3 nm
FRC XZ resolution = 47.1 nm
FRC YZ resolution = 47.6 nm
Cell 2 LS
Whole Slice
660,758 localizations
FRC XY resolution = 42.0 nm
FRC XZ resolution = 53.8 nm
FRC YZ resolution = 48.6 nm
Cell 3 LS
Whole Slice
637,511 localizations
FRC XY resolution = 40.3 nm
FRC XZ resolution = 45.9 nm
FRC YZ resolution = 45.0 nm
Fourier Ring Correlation
Spatial Frequency (nm⁻¹)
0
0.02
0.04
0.06
1

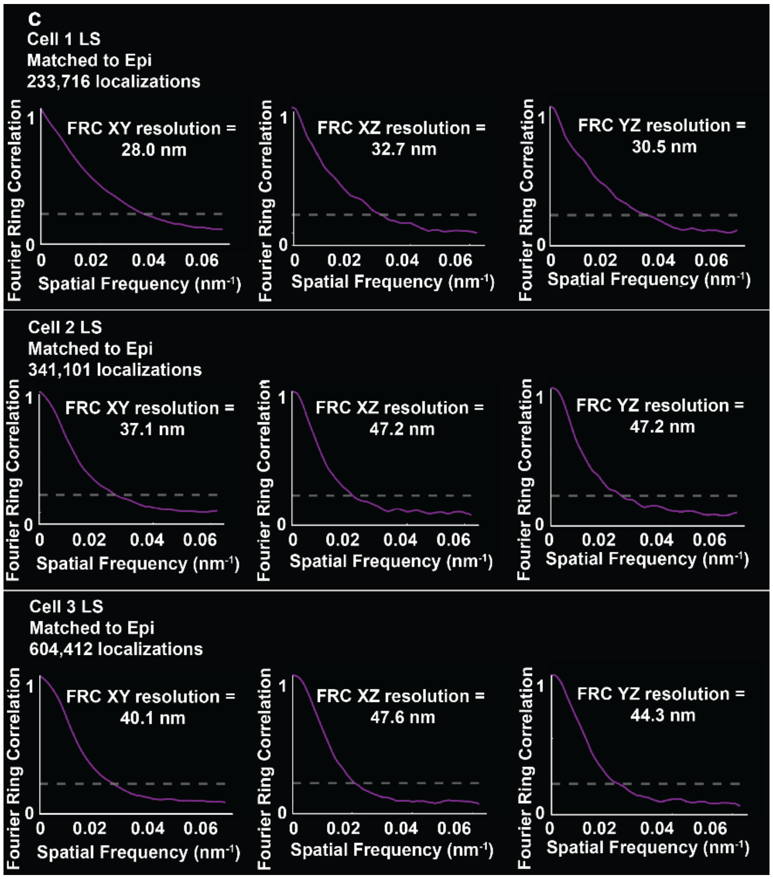


**Supplementary Fig. 10. Fourier ring correlation (FRC) curves for light sheet (LS) compared to epi-illumination for 3D high-density single-molecule super-resolution imaging of lamin A/C. a** FRC in the xy, xz, and yz planes, respectively, for the reconstruction of a whole 3D slice imaged with epi-illumination for three different cells. The FRC resolutions in xy/xz/yz are 41.3/59.7/54.2 nm from 233,716 localizations for cell 1, 42.3/56.6/50.3 nm from 341,101 localizations for cell 2, and 42.2/51.0/46.4 nm from 604,412 localizations for cell 3. **b** FRC in the xy, xz, and yz planes, respectively, for the reconstruction of the same whole 3D slice imaged with LS illumination for the same three cells as in a. The FRC resolutions in xy/xz/yz are 37.3/47.1/47.6 nm from 1,215,297 localizations for cell 1, 42.0/53.8/48.6 nm from 660,758 localizations for cell 2, and 40.3/45.9/45.0 nm from 637,511 localizations for cell 3, demonstrating improved number of localizations and resolution for LS compared to epi-illumination. **c** FRC in the xy, xz, and yz planes, respectively, for the reconstruction of a whole 3D slice imaged with LS illumination after the localizations with the best localization precisions are kept to match the number of localizations acquired with epi-illumination for the three different cells in a. The FRC resolutions in xy/xz/yz are 28.0/32.7/30.5 nm for cell 1, 37.1/47.2/47.2 nm for cell 2, and 40.1/47.6/44.3 nm for cell 3, demonstrating improved resolution over epi-illumination even when the numbers of localizations are matched. All data was filtered for localization precision less than 30 nm in xy. A high imager strand concentration of 0.1 nM was used for all single-molecule acquisitions. 10,000 frames were acquired for each data set and analyzed using DECODE.

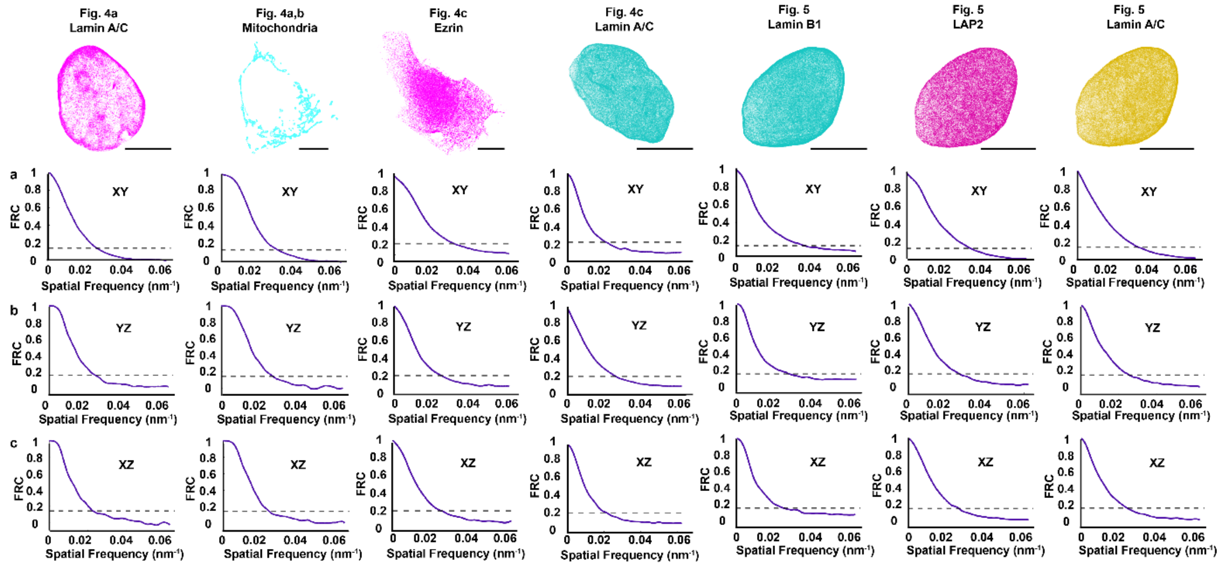


**Supplementary Fig. 11. Statistics for double-helix PSF localization data used for the 3D super-resolution reconstructions of lamin A/C, mitochondria, and ezrin in Fig. 4 and of lamin B1, LAP2, and lamin A/C in Fig. 5. a** Fourier ring correlation (FRC) for the xy plane, **b** FRC for the yz plane, and **c** FRC for the xz plane. The FRC resolutions in xy/yz/xz for the reconstructions in Fig. 4a,b were 35.0/45.7/51.5 nm for lamin A/C and 32.7/40.2/44.9 nm for mitochondria. The FRC resolutions in xy/yz/xz for the reconstructions in Fig. 4c were 31.0/39.2/38.2 nm for ezrin and 38.0/46.3/47.2 nm for lamin A/C. The FRC resolutions in xy/yz/xz for the reconstructions in Fig. 5 were 27.9/36.1/39.9 nm for lamin B1, 30.2/37.6/38.0 nm for LAP2, and 31.3/38.9/40.6 nm for lamin A/C. Scale bars 10 μm.

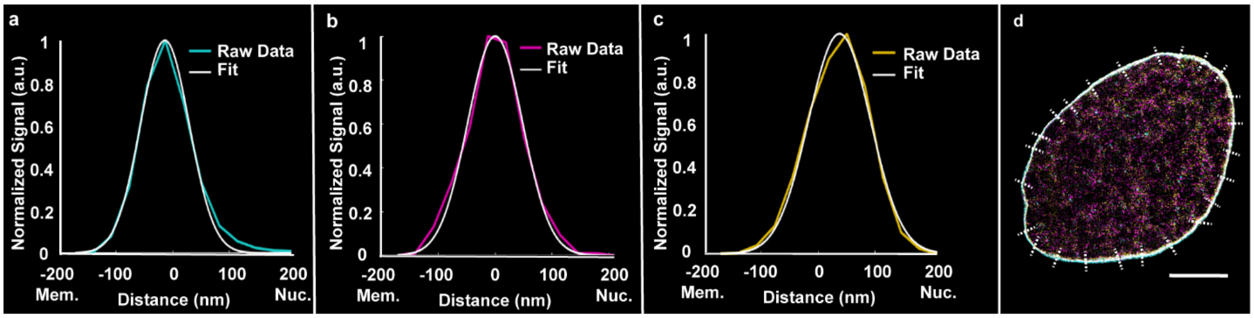


**Supplementary Fig. 12. Representative line scans and statistical analysis of distance determinations for lamin B1, LAP2, and lamin A/C.** Example line scan data and corresponding Gaussian fits for line scans and distance plots in Fig. 5b shown for **a** lamin B1, **b** LAP2, and **c** lamin A/C. **d** A total of 25 line scans were acquired along the nuclear rim of a 500 nm-thin slice of all three targets. Scale bar 10 μm.

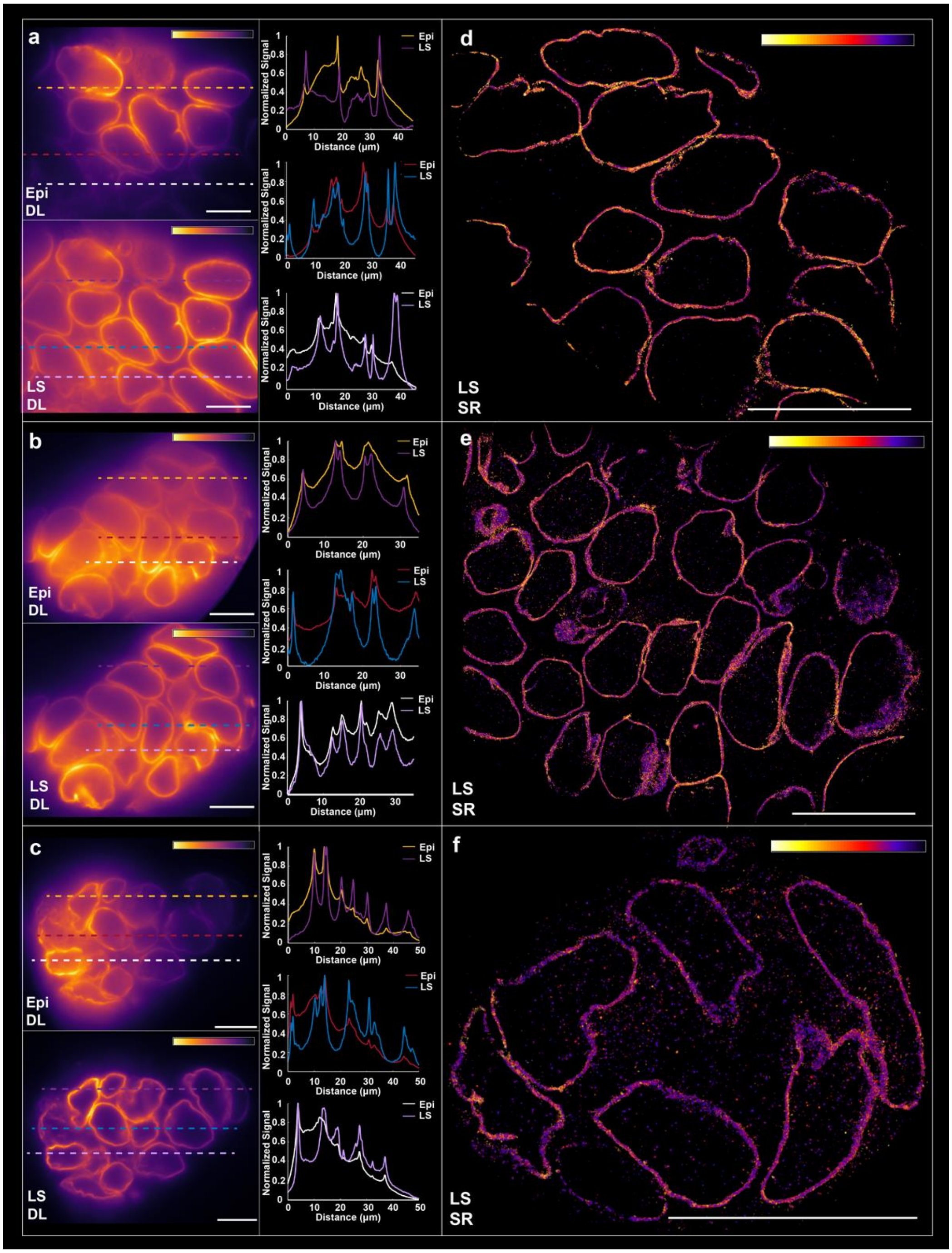


**Supplementary Fig. 13. Diffraction-limited (DL) and super-resolution (SR) imaging of stem cell aggregates. a-c** Line scans across three different stem cell aggregates cultured in the microfluidic chips and imaged with epi-illumination (top) and light sheet (LS) illumination (bottom), demonstrating the consistent background reduction capabilities of soTILT3D, even when imaging larger samples. Scale bars 10 μm. **d-f** Single-molecule super-resolution imaging of three different stem cell aggregates cultured in the microfluidic chips and imaged with LS illumination demonstrating the versatility of soTILT3D for imaging diverse biological samples and the adaptability of the microfluidic device to enable sample culturing and imaging across different scales. Scale bars 20 μm. The colorbars show intensity normalized independently for each image.

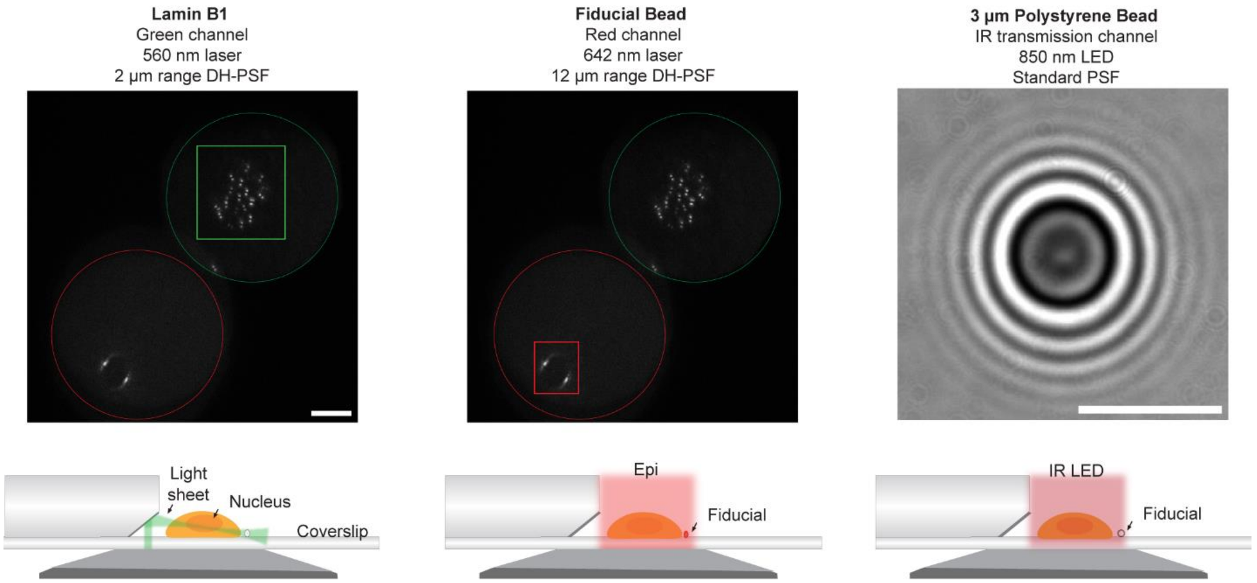


**Supplementary Fig. 14. Illumination and detection scheme for the soTILT3D platform.** Labeled cellular structures were excited by 560 nm single-objective light sheet illumination and detected in the green channel using the 2-µm axial range double-helix (DH) PSF. Fluorescent fiducial beads adhered to the coverslip were excited by 642 nm epi-illumination and detected in the red emission channel using the 12-µm axial range DH-PSF. 3 µm polystyrene beads adhered to the coverslip were imaged using transmitted IR light detected by a separate CMOS camera. The diffraction rings were used to perform active stabilization of the sample during long-term imaging. Scale bars 10 µm.

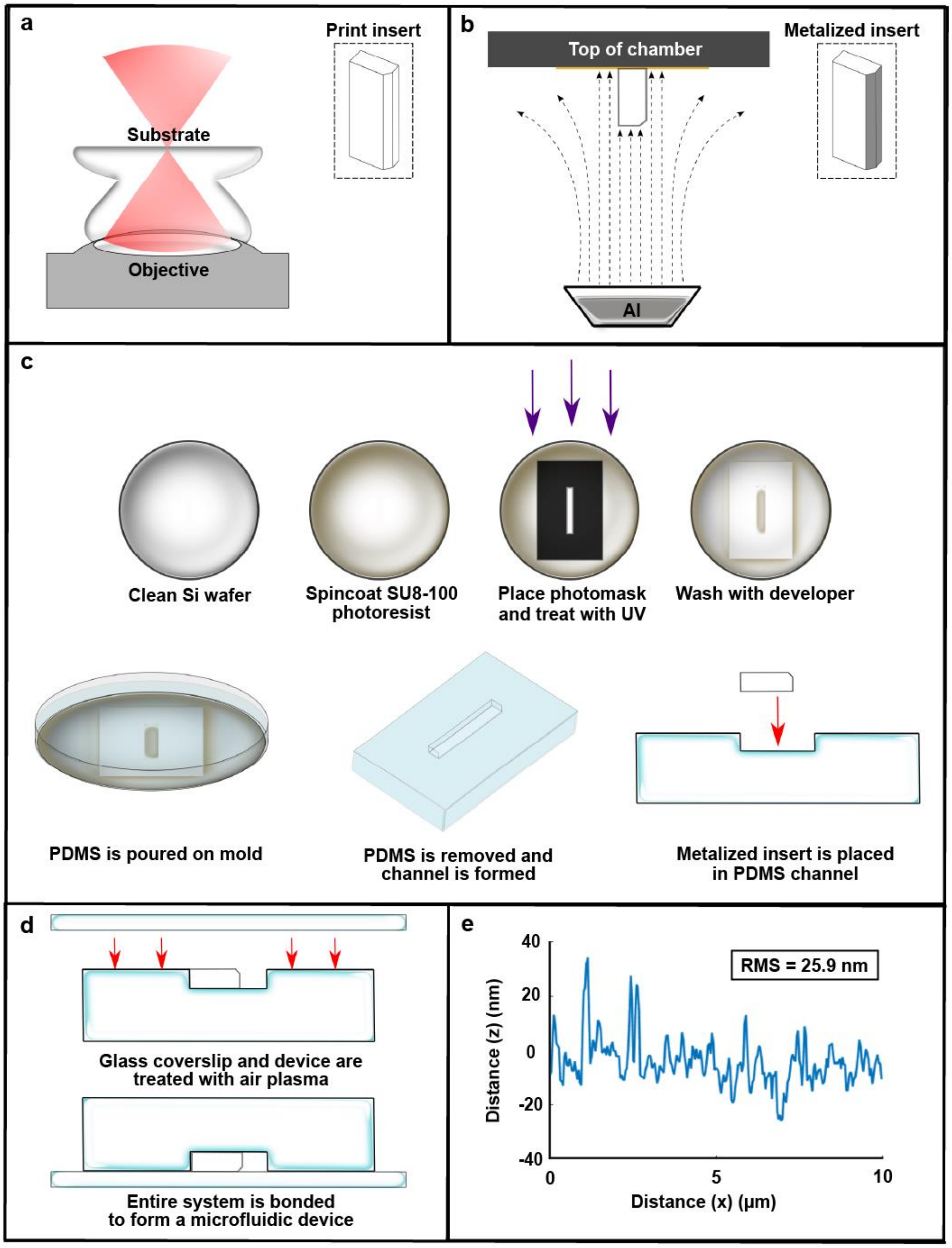


**Supplementary Fig. 15. Schematic of the microfluidic chip fabrication pipeline. a** Illustration of the two-photon polymerization approach utilized for nanoprinting the microfluidic insert. **b** Schematic of metal vapor deposition setup used on the side walls of the printed insert. **c** Protocol used to fabricate channels out of PDMS using soft lithography with molds made from SU8-100 photoresist on Si wafers along with schematic of insert being positioned inside of a PDMS channel. **d** Schematic showing PDMS being bonded to a glass coverslip to form a microfluidic device. **e** Representative example of a 10 μm line profile along the surface of the metalized side wall using atomic force microscopy (AFM). The roughness root mean square of 10 $\mu m^2$ of the surface was determined to be 25.9 nm.

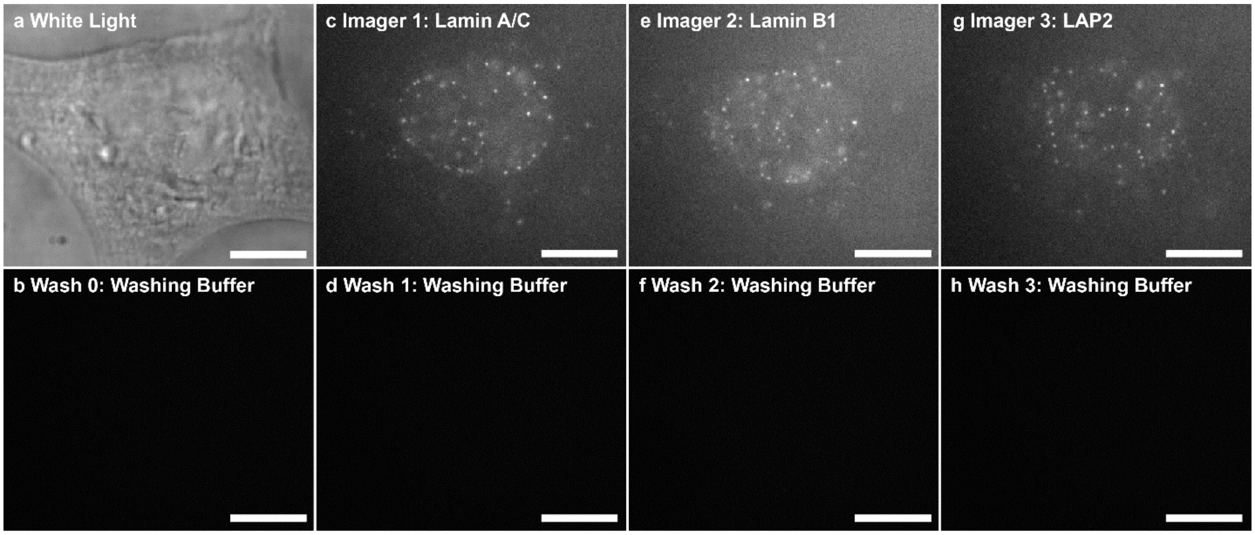


**Supplementary Fig. 16. DNA-PAINT imaging controls.** 1,000 frames for each target were acquired in 2D with the same imaging settings and analyzed in ThunderSTORM with the same settings and filters. **a** White light image of a cell labeled for lamin A/C, lamin B1, and LAP2 and **b** wash 0 with washing buffer which yielded 33 localizations. Single-molecule data **c** of 0.1 nM imager strand concentration for lamin A/C, which yielded 27,838 localizations, **d** after wash 1 with washing buffer, which yielded 750 localizations, **e** of 0.04 nM imager strand concentration for lamin B1, which yielded 22,276 localizations, **f** after wash 2 with washing buffer, which yielded 639 localizations, **g** of 0.1 nM imager strand concentration for LAP2, which yielded 27,028 localizations, and **h** after wash 3 with washing buffer, which yielded 575 localizations. Scale bars 10 µm.

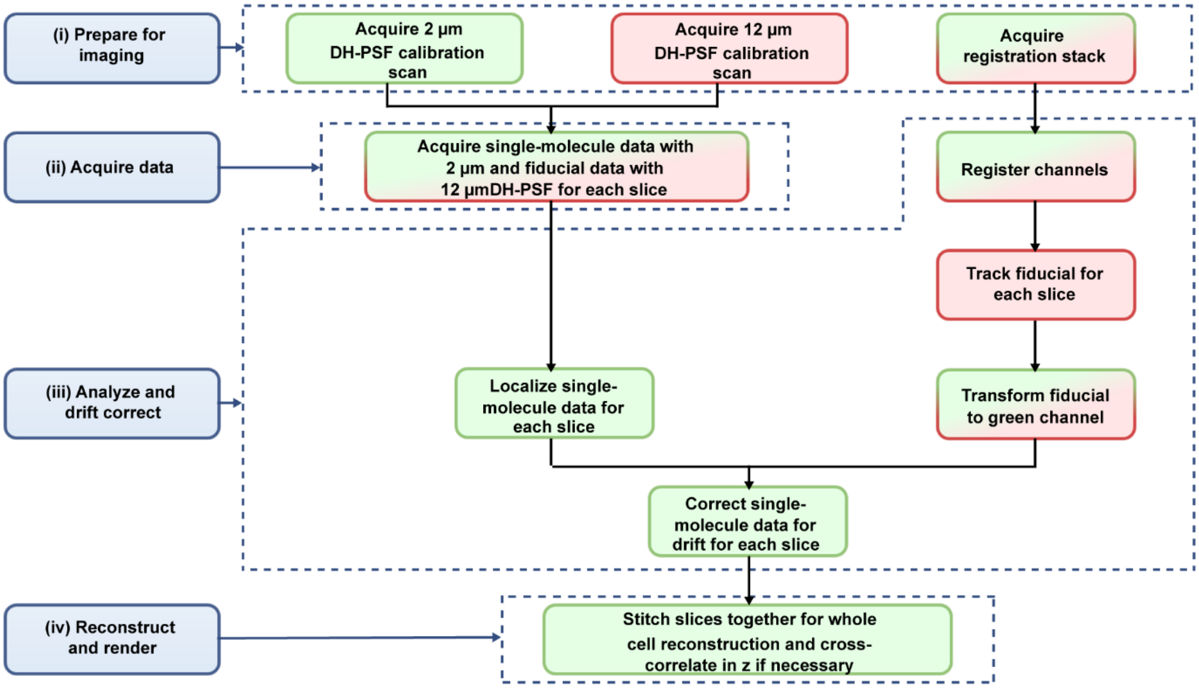


**Supplementary Fig. 17. Imaging and analysis workflow.** Flowchart showing the steps for imaging and analysis for the reconstructions shown in Figs. 4 and 5.

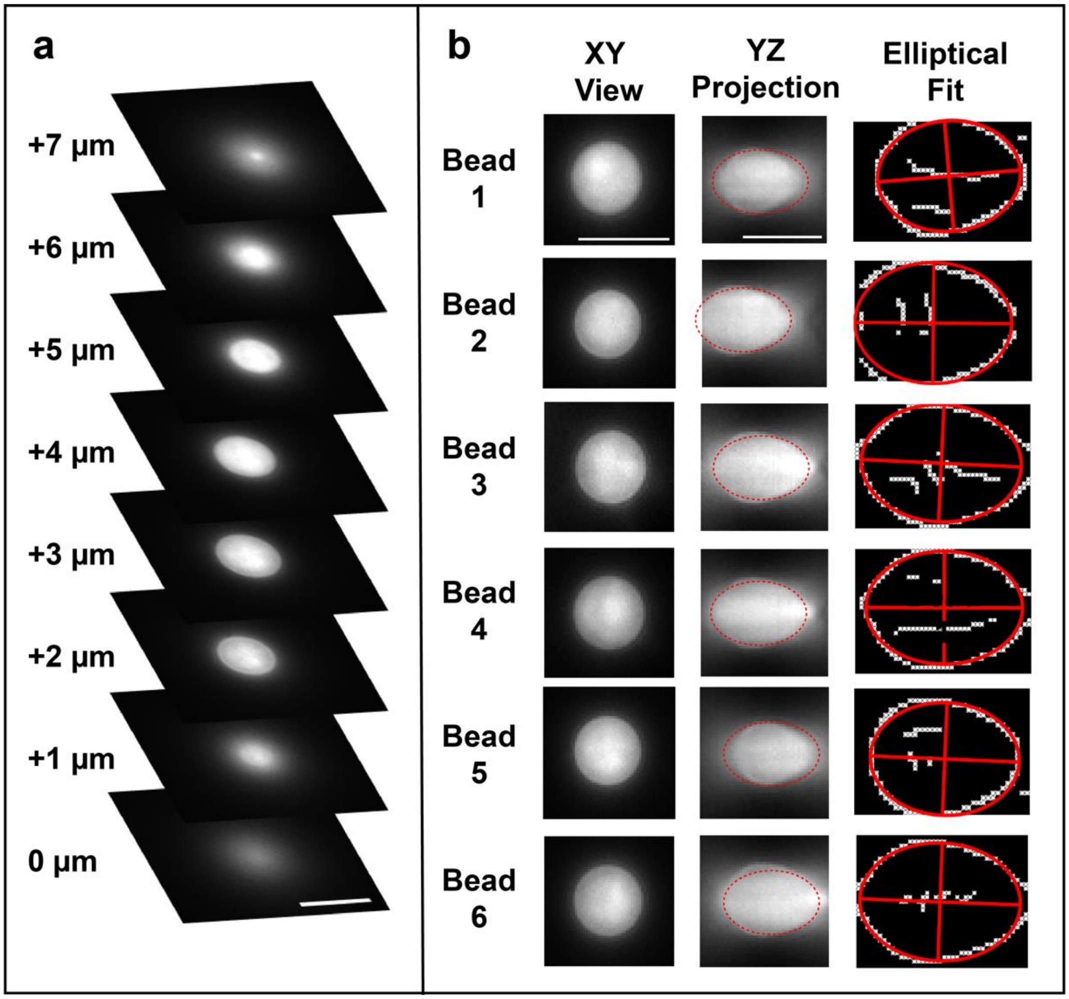


**Supplementary Fig. 18. Empirical measurements of the axial compression factor. a** Example images of a z stack of Bead 4 in b. In total, six separate 4 µm beads were imaged. Scale bar 5 µm. **b** An ellipse was fit to the YZ projection of each bead and the axial compression factor was determined by taking the ratio of the minor and major axes of the fitted ellipse. This yielded a compression factor of 0.75 ± 0.02 (mean ± standard deviation for the six beads). Scale bars 5 µm.

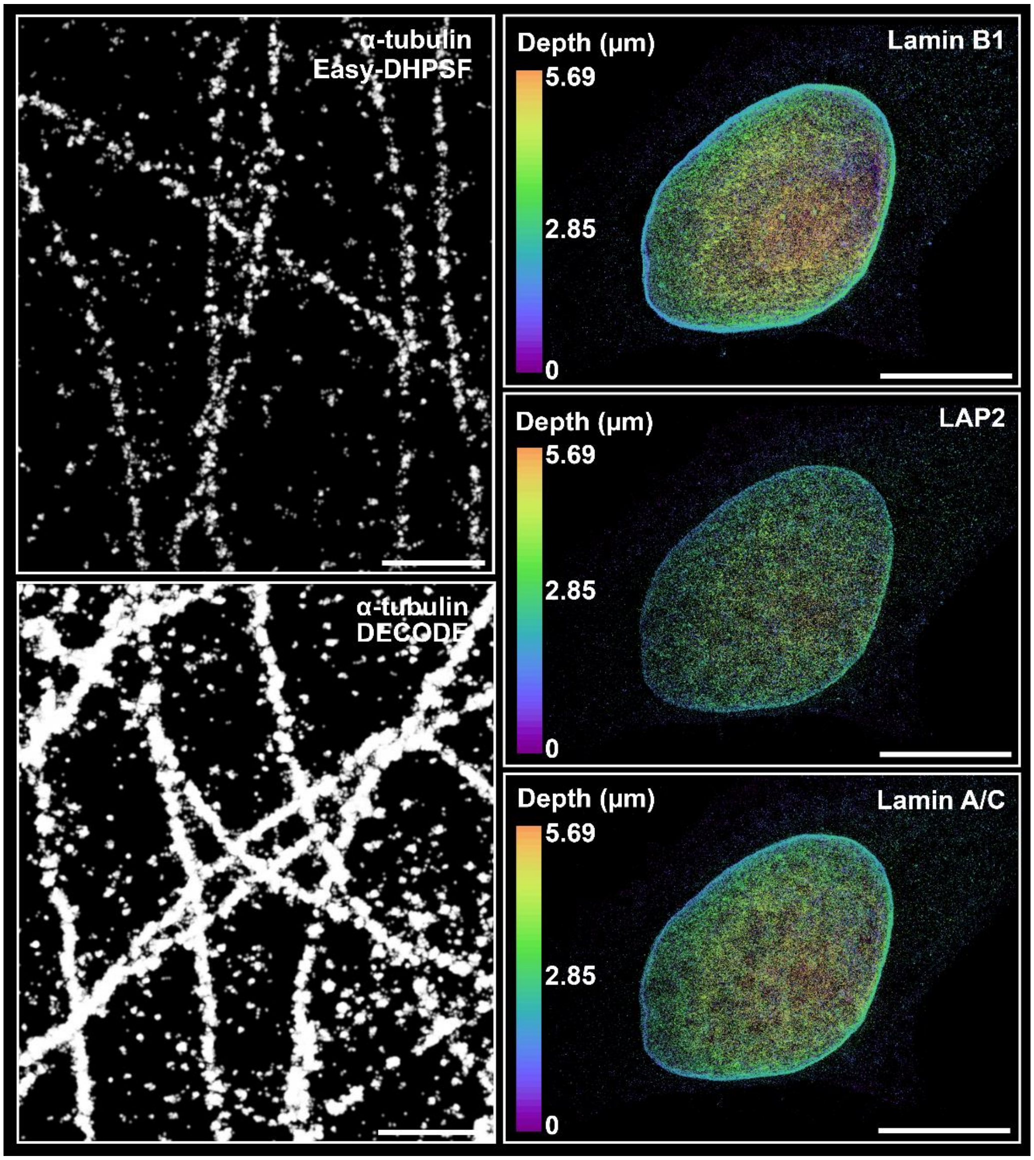


**Supplementary Fig. 19. Reconstructions for double-helix PSF localization data used for the 3D super-resolution reconstructions of microtubules in Fig. 3a and of lamin B1, LAP2, and lamin A/C in Fig. 5 without filtering of spurious localizations.** The FRC resolutions in xy/yz/xz for the reconstructions of microtubules in Fig. 3a when including spurious localizations were 36.1/41.8/42.2 nm for the Easy-DHPSF analyzed sample and 29.8/37.2/42.2 nm for the DECODE analyzed sample. Scale bars 1 μm. The FRC resolutions in xy/yz/xz nm for the reconstructions in Fig. 5 when including spurious localizations were 22.5/29.7/29.0 nm for lamin B1, 31.8/39.5/41.4 nm for LAP2, and 32.7/40.2/43.6 nm for lamin A/C. Scale bars 10 μm.

**Supplementary Table 1. Acquisition and reconstruction times for 3D super-resolution (SR) reconstructions.** Acquisition and reconstruction times for all 3D SR reconstructions. Fitting for reconstruction was performed using a DECODE model that took ~12 hours to train. The exposure time was 100 ms for all targets.

| | **Number of frames per slice** | **Number of slices** | **Total acquisition time (minutes)** | **Reconstruction time (minutes)** |
|---|---|---|---|---|
| **Fig. 4a,b Lamin A/C** | 50,000 | 1 | ~80 | ~25-50 |
| **Fig. 4a,b Mitochondria** | 50,000 | 1 | ~80 | ~25-50 |
| **Fig. 4c Lamin A/C** | 10,000 | 4 | ~70 | ~20-40 |
| **Fig. 4c Ezrin** | 20,000 | 4 | ~130 | ~40-80 |
| **Fig. 5 Lamin B1** | 10,000 | 8 | ~130 | ~40-80 |
| **Fig. 5 Lamin A/C** | 10,000 | 8 | ~130 | ~40-80 |
| **Fig. 5 LAP2** | 10,000 | 8 | ~130 | ~40-80 |
